# Blind Recognition Reveals Early Multiphase IL-1ra Aggregation

**DOI:** 10.64898/2026.08.03.742394

**Authors:** Andrei A. Raibekas

## Abstract

The primary question in this work is whether a turbidity trace contains more than one resolved kinetic event before any rate or aggregate-mass information is supplied. A blind, three-gate hierarchical Gompertz procedure applied to human interleukin-1 receptor antagonist (IL-1ra) at 53 °C in phosphate retained one phase at 1 and 2 mg/mL, but selected two phases at 5 and 10 mg/mL. Thus the first resolved multiphase behavior appears at the low protein concentration of 5 mg/mL. Source optical-rate markers were withheld until after selection. Those observed optical rates were then compared with blind-fit phase rates, and the same comparison was expressed on a conditional mass-equivalent scale using the published 5.3-fold Type-I/Type-II turbidity response ratio. A separate 50 °C citrate series (4, 6, 8, 10, and 14 mg/mL) also selected two phases in every retained trace; its printed Vmax values were used only as post-selection comparators. Finally, two challenging 40 °C high-concentration traces (180 and 200 mg/mL) were digitized from the instrument-output plot and each retained two phases. All three conditions are analyzed independently. Blind recognition is therefore the central result; the mass-equivalent conversion is a secondary, post-selection interpretation of optical phase rates. The absolute *µ*M*/*min scale remains conditional on an endpoint aggregate fraction measured elsewhere.

## 1 Introduction

Turbidity is a useful kinetic readout, but a visually smooth rise need not be one kinetic event. A whole-trace sigmoid can average over sequential or overlapping aggregate populations, and an optical signal is not generally proportional to aggregate mass because particle properties affect scattering [1, 2, 3]. In conventional Gompertz analysis, a single curve is often fitted to the full trace, while decisions to add a second component may depend on visual inspection, initial guesses, or manual trial and error. Such practice can conceal an early low-amplitude event or turn broad overlap into an unsupported extra phase.

Blind recognition is defined as selecting the number of kinetic phases from the time course alone, before observed rate markers, aggregate-mass information, or the published turbidity response factor are introduced. The procedure starts with *K* = 1 and tests additional ordered Gompertz components. A new component is accepted only if three prespecified gates are passed: (i) the improvement is substantial, ΔAICc < − 10; (ii) each component amplitude exceeds 10% of the total fitted amplitude; and (iii) each adjacent inflection-time gap exceeds twice the preceding component’s time constant. Thus, blind recognition is blind to external rate and mass information, not to the time course or to the stated model family.

Compared with normally applied manual Gompertz fitting, this hierarchy makes the phase-count decision reproducible and auditable. It reduces dependence on visual judgment and starting guesses, limits unsupported over-splitting, and applies the same rule to early, overlapping, and visually obscured traces. The method is therefore a conservative recognition tool, not a claim of a new microscopic mechanism or a universal kinetic law.

The work has two primary, deliberately separate rate-comparison series. The 53 °C phosphate series establishes where blind recognition first resolves multiphase behavior, including the low-concentration range up to 5 mg/mL. The 50 °C citrate series is analyzed by the same blind framework as a separate condition-specific analysis. A third 40 °C high-concentration series is a deliberately demanding recognition test: its visually obscured late behavior is assessed without observed-rate or mass conversion. The conditions differ in formulation, temperature, and concentration range, so their absolute rates are not normalized or pooled.

## 2 Data and methods

### 2.1 Protein production and aggregation-kinetics measurement

The recombinant human interleukin-1 receptor antagonist (IL-1ra) was expressed in *Escherichia coli*, refolded from inclusion bodies, and purified by column chromatography, as previously described [4]. The aggregation kinetics of IL-1ra were measured using a 96-well glass-bottom plate and a temperature-controlled plate reader (SpectraMax Plus, Molecular Devices, Sunnyvale, CA) [5, 6]. The sample volume was 0.2 mL per well. The plates were incubated in the spectrophotometer at temperatures between 40 and 53 °C, and the optical density was measured at 450 nm at intervals of either 1 or 2 min, with an automated shaking step of 3 s between each reading.

### 2.2 Three condition-specific kinetic datasets

The 53 °C phosphate series comprises OD_450_ traces at 1, 2, 5, and 10 mg/mL in 10 mM sodium phosphate, pH 6.5, 140 mM NaCl, and 0.5 mM EDTA [6]. Each trace contains 46 approximately one-minute samples. Source optical-rate markers of 104.6, 237.4, 408.0, and 552.5 mOD/min were withheld from fitting and phase selection.

The 50 °C citrate series comprises traces at 4, 6, 8, 10, and 14 mg/mL in 10 mM citrate, pH 6.5, 140 mM NaCl, and 0.5 mM EDTA. The 12 mg/mL trace is excluded throughout for clarity. These traces were digitized at 50-s spacing from 0 to 7000 s. Their printed moving-window Vmax values are retained only as observed optical comparators after blind selection; they are not inputs to fitting, phase selection, or conversion-factor estimation.

The 40 °C high-concentration challenge set comprises the illustrative 180 and 200 mg/mL traces. It is retained as a separate condition and is not used for cross-series kinetic or mass-rate comparison. The available record is a high-resolution SigmaPlot-style graphic derived from the instrument output rather than the native numerical export. Marker coordinates were digitized on the 50-s display grid; the 180 mg/mL trace contained a visible positive instrument offset at approximately 5350 s. A 0.028 OD post-offset correction and one straight-line bridging point were applied before fitting, with each reconstructed value and correction flag retained in the accompanying data file.

The 53 °C and 50 °C primary series share pH 6.5, 140 mM NaCl, and 0.5 mM EDTA. The sole formulation-buffer difference is 10 mM phosphate in the 53 °C series versus 10 mM citrate in the 50 °C series. Temperature (53 °C versus 50 °C) and protein-concentration range (1–10 versus 4–14 mg/mL) also differ. These differences require within-series interpretation; the separate 40 °C challenge set is not pooled with either series.

### 2.3 Blind hierarchical phase recognition

The nested optical model is

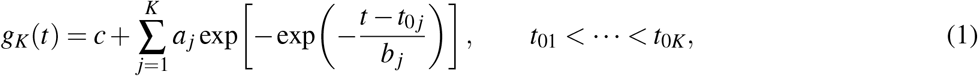

where the peak optical rate of component *j* is

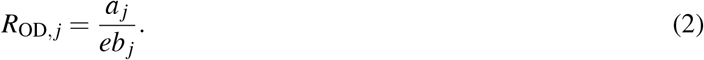

Model comparison used the Akaike Information Criterion corrected for small sample size (AICc), which scores a fitted model by combining its goodness of fit with a penalty for the number of free parameters [10]. Starting at *K* = 1, an added component was accepted only when ΔAICc *< −*10, every amplitude exceeded 10% of the total fitted amplitude, and every adjacent inflection-time gap exceeded two preceding-component time constants. In short,the recognition path is linear and stops at the first failure: *K* = 1 → fit *→*test *K* + 1 against the three gates above *→*if all three pass, set *K* = *K* + 1 and repeat; if any one fails, stop and retain the current *K →*continue until *K*_max_ is reached or a gate fails. The 53 °C and 40 °C series used *K*_max_ = 2. The 50 °C series used *K*_max_ = 3, ten independent starts, and at least 80% recurrence of the selected phase count. No observed rate, mass estimate, or response factor was supplied to this procedure.

### 2.4 Operating-characteristic and sensitivity checks

To test the behavior of the hierarchy when the ground truth is known, four synthetic traces were generated using the same ordered Gompertz model: a one-phase trace, a clearly separated two-phase trace, a strongly overlapping two-phase trace, and a two-phase trace whose second amplitude was below the 10% acceptance threshold. Each trace contained 131 points at 50-s spacing and independent Gaussian optical noise with standard deviation 0.008 OD. One hundred independent realizations of each case were fitted with the same model hierarchy and gates. These simulations are operating-characteristic checks, not independent biological validation.

Sensitivity to the digitized 40 °C records was also tested by adding independent Gaussian perturbations of standard deviation 0.010 OD to every reconstructed point and repeating the complete one-versus-two-phase fit 100 times for each trace. To test the 40 °C result against a matched model-based null, 100 one-phase null traces were generated from each trace’s fitted *K* = 1 curve and 100 two-phase alternatives from its fitted *K* = 2 curve, preserving the observed time points and adding the same 0.010 OD noise. For the 180 mg/mL trace, the post-offset correction was additionally drawn from 0.0282*±*0.0050 OD and the analysis was repeated with the bridge point retained or omitted. These are operating-characteristic and sensitivity checks within the Gompertz model family; they are not measurement-error estimates, tests of model misspecification, or independent biological validation. The random seed, simulation summary, and analysis scripts are included in the accompanying package.

### 2.5 Post-selection conversion to a mass-equivalent rate

Krishnan and Raibekas measured isolated Type I and Type II IL-1ra aggregates and reported a 5.304-fold higher OD_450_ response per unit mass for Type I material (their Fig. 1B) [5]. Their aggregate-mass measurements motivate the conditional endpoint aggregate fraction *f*_agg_ = 0.22 used here. No source-paper turbidity-versus-mass plot is reproduced, and neither the response ratio nor endpoint fraction enters phase selection.

**Figure 1:**
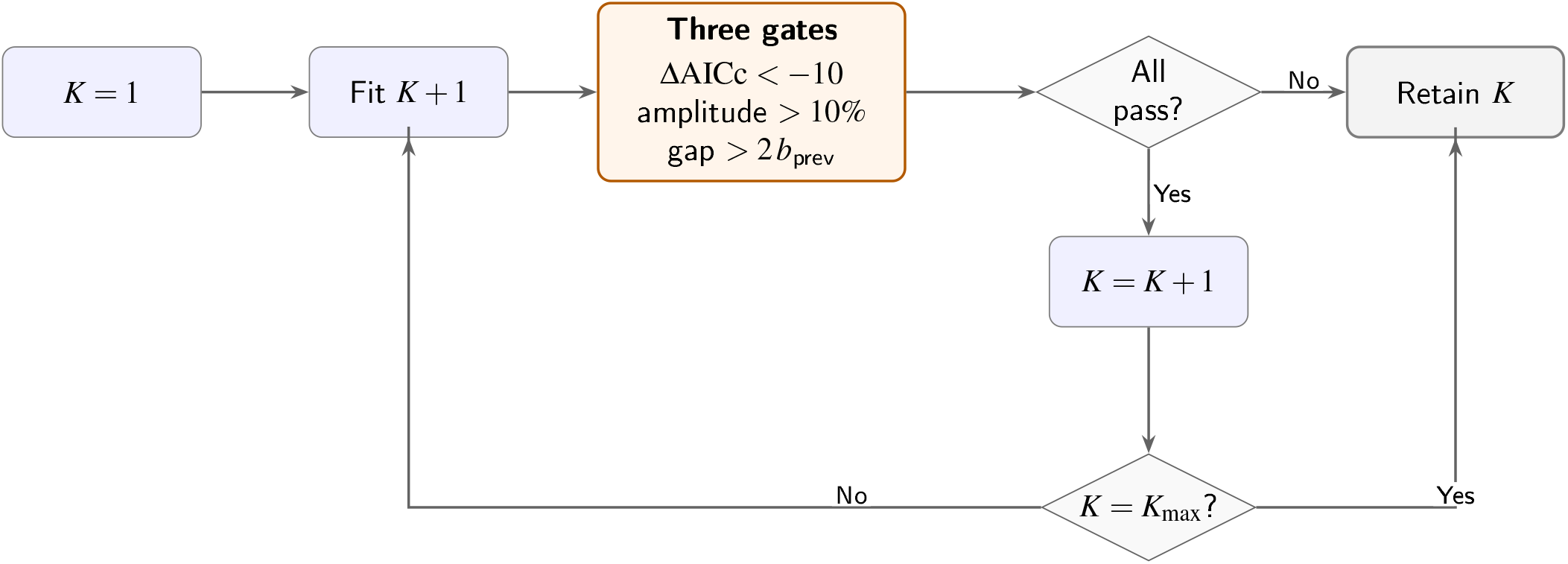
Blind hierarchical phase-recognition procedure. The loop starts by testing a two-phase fit against the one-phase baseline using the three gates. If any gate fails, one phase is retained and the procedure stops. If all three pass, two phases become the accepted result; the procedure then stops at two phases if *K*_max_ has been reached, or otherwise tests a three-phase fit against the two-phase baseline using the same gates, retaining three phases only if that fit also passes.

**Figure 2:**
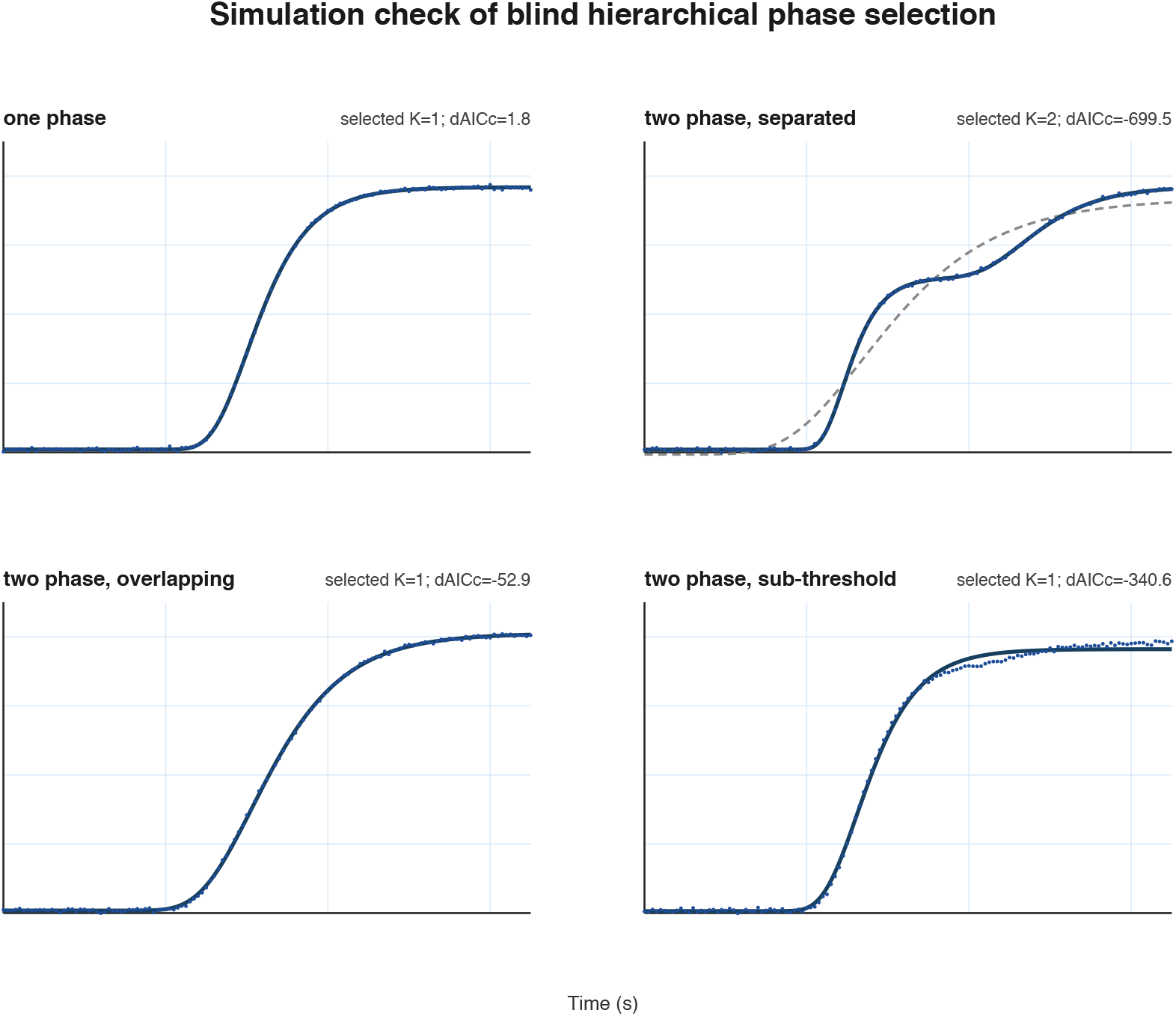
Simulation check of the blind hierarchy. Each panel shows one noisy realization with its one-phase fit (grey dashed) and the selected model (navy). The clearly separated two-phase trace is retained, whereas strongly overlapping and sub-threshold two-phase traces are conservatively retained as one phase. These simulations test operating characteristics and do not establish biological mechanism.

Assigning phase 1 to the high-response population and phase 2 to the low-response population gives *α*_1_*/α*_2_ = 5.304. For a two-phase trace, terminal mass-equivalent fractions are

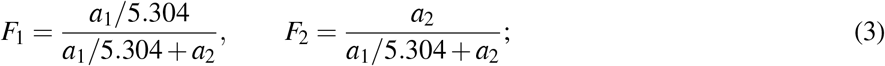

for a selected one-phase trace, *F*_1_ = 1. With molecular weight 17.3 kDa, total protein concentration is *C*_tot_ =57.803*p µ*M for *p* in mg/mL. The conditional phase rate is

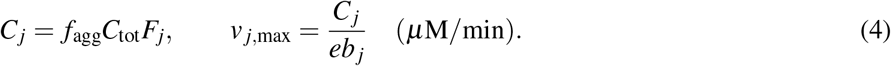

The 0.2-mL sample volume converts a concentration rate to amount rate by *n*?_*j*_ = 0.2*v*_*j*_ nmol/min, but does not change *µ*M*/*min.

For the observed comparator, its optical rate is multiplied by the blind phase-1 conversion *v*_1,max_*/R*_OD,1_ for the same trace. The result is explicitly termed an *observed phase-1-equivalent µ*M*/*min value. It is not a direct independently measured mass rate. This transformation lets the observed-versus-fit difference be viewed both before and after conversion without claiming phase-specific mass data for the moving-window Vmax.

## 3 Results

### 3.1 Synthetic operating-characteristic checks

The simulation results define both the useful operating range and the conservative boundary of the procedure. Under the stated noise level, the hierarchy retained one phase in all one-phase realizations and retained two phases in all clearly separated two-phase realizations. In the strongly overlapping two-phase case, only 4% of realizations passed all gates; in the sub-threshold case, only 5% passed. In both difficult cases, AICc alone favored two components, but the structural gates suppressed a split that was not reliably identifiable from the whole trace. This behavior is intentional and is the reason the method should be described as phase recognition with conservative acceptance, not automatic phase enumeration.

For the 40 °C challenge specifically, the matched one-phase null selected *K* = 2 in 0/100 runs at both 180 and 200 mg/mL, while matched two-phase alternatives were recovered in 100/100 and 99/100 runs, respectively. The direct digitization perturbation retained *K* = 2 in 100/100 runs for both traces. For the 180 mg/mL trace, varying the offset correction and retaining or omitting the bridge point also retained *K* = 2 in 100/100 runs. These results support stability of the selected phase count to the tested digitization and correction uncertainties, while not replacing validation against native instrument exports or ruling out broader model misspecification.

### 3.2 53 °C phosphate series

#### 3.2.1 Blind recognition resolves multiphase behavior at 5 mg/mL

One-phase fits appear visually strong at all four concentrations, but the blind hierarchy retained *K* = 1 at 1 and 2 mg/mL and selected *K* = 2 at 5 and 10 mg/mL (Fig. 3). At 5 and 10 mg/mL, the two-phase improvement was large (ΔAICc = -173.2 and -154.7, respectively). The candidate split at 1 mg/mL failed temporal separation, while that at 2 mg/mL lacked information support. The first resolved multiphase record is therefore 5 mg/mL, before the concentration range becomes high.

**Figure 3:**
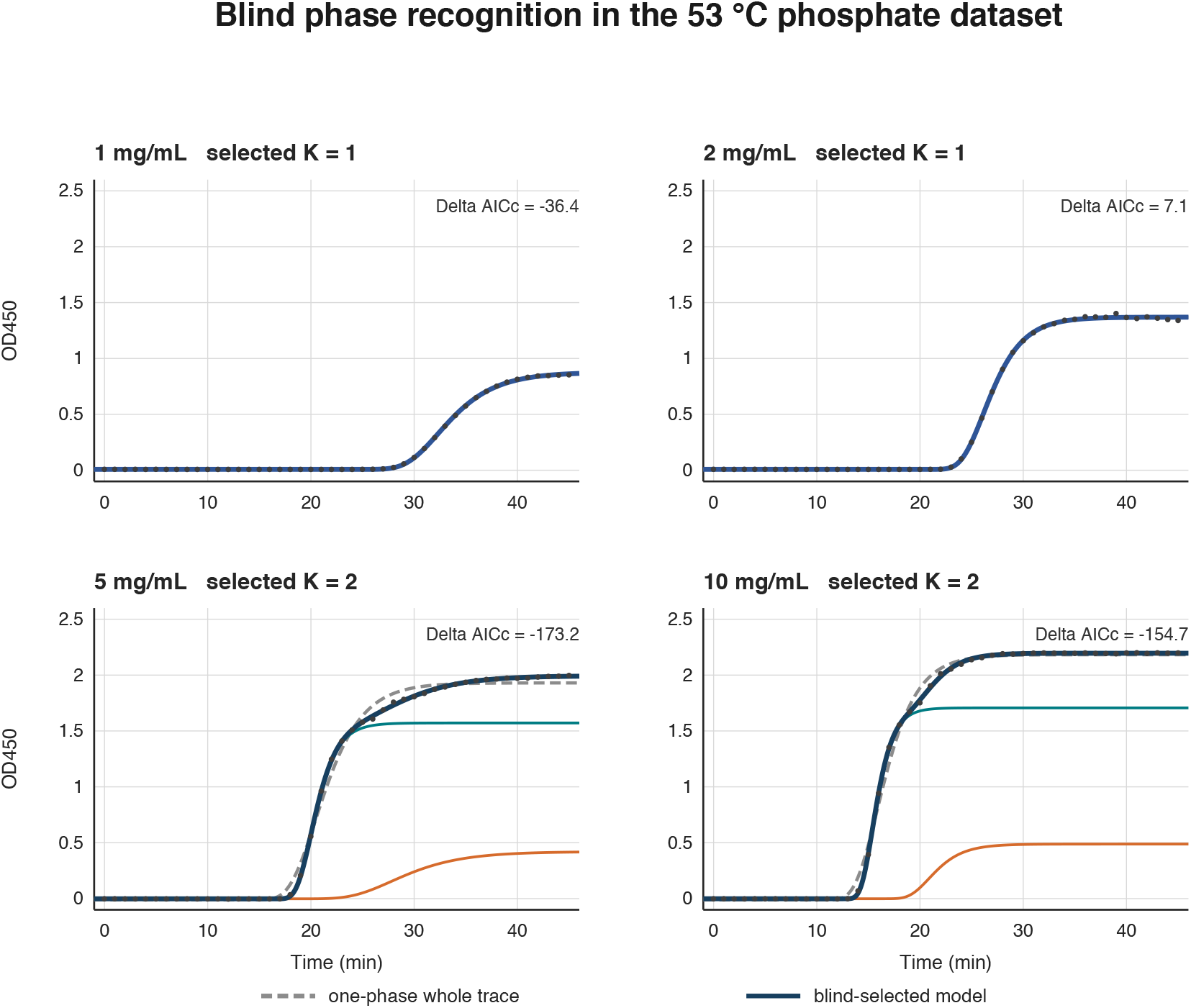
Blind phase recognition in the 53 °C phosphate series. Grey dashed curves are one-phase whole-trace fits; navy curves are blind-selected totals; teal and orange curves are the selected component contributions. The 1 and 2 mg/mL records remain one-phase; the 5 and 10 mg/mL records select two phases.

#### 3.2.2 Observed and blind-fit rates agree for the initial event

The withheld source markers were consulted only after phase selection. Table 2 compares each observed optical rate with the blind phase-1 rate and, where present, the blind phase-2 rate. It repeats the comparison after post-selection conversion to phase-1-equivalent *µ*M*/*min. The source markers are close to the blind phase-1 rates at all concentrations; their signed differences are -0.7, -1.3, +0.9, and +2.0%, respectively. The phase-2 rates are reported as additional blind predictions, not as a comparison against an observed phase-2 marker.

**Table 1:**
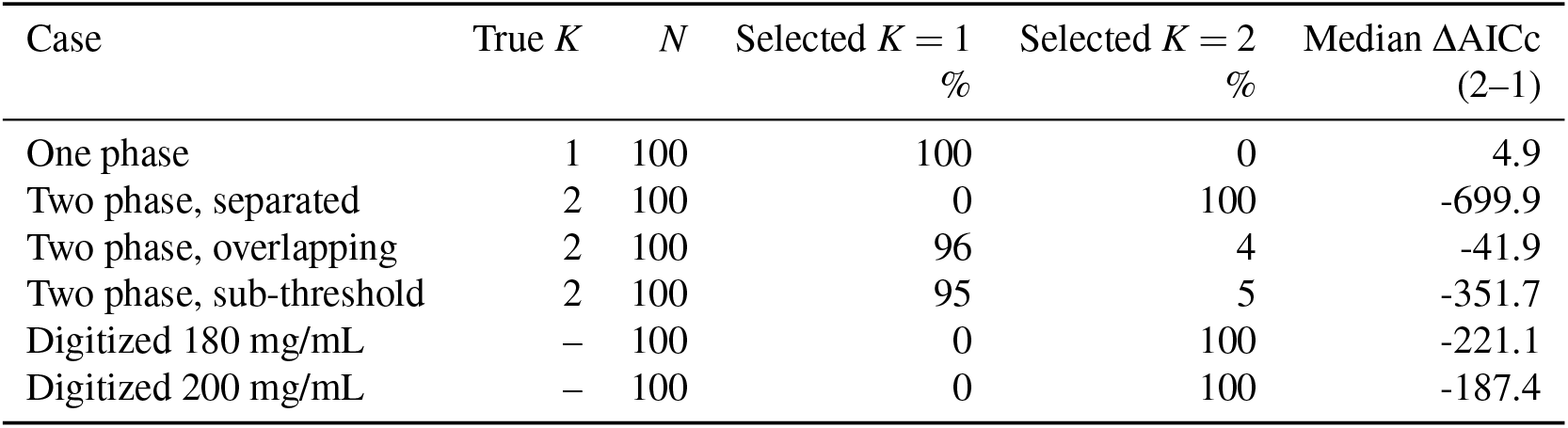
Synthetic and perturbation checks. Percentages are the fraction of 100 complete refits selecting the indicated phase count. Synthetic traces used 0.008 OD noise; digitized traces used 0.010 OD pointwise perturbations.

**Table 2:**
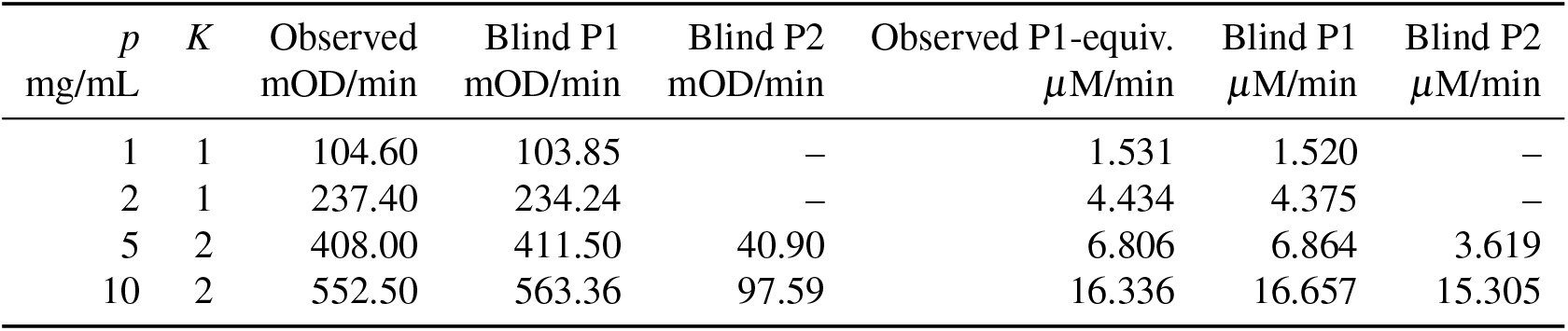
53 °C phosphate series: observed versus blind-fit rates. “Observed” is the withheld source optical marker. Observed phase-1-equivalent mass rates use the same-trace blind phase-1 optical-to-mass scaling; all *µ*M*/*min values assume *f*_agg_ = 0.22.

### 3.3 50 °C citrate series

#### 3.3.1 Blind recognition is two-phase throughout the retained range

The 50 °C citrate records are analyzed separately, with no transfer of the 53 °C phase count, rates, or calibration. The blind hierarchy selected *K* = 2 at 4, 6, 8, 10, and 14 mg/mL (Fig. 4). The selected phase count recurred in all ten starts at 4, 6, and 8 mg/mL, and in eight of ten starts at 10 and 14 mg/mL. Candidate third components were rejected as too small or unresolved. The compact redraw makes the early transition and later increment legible without the elongation of the original slide display.

**Figure 4:**
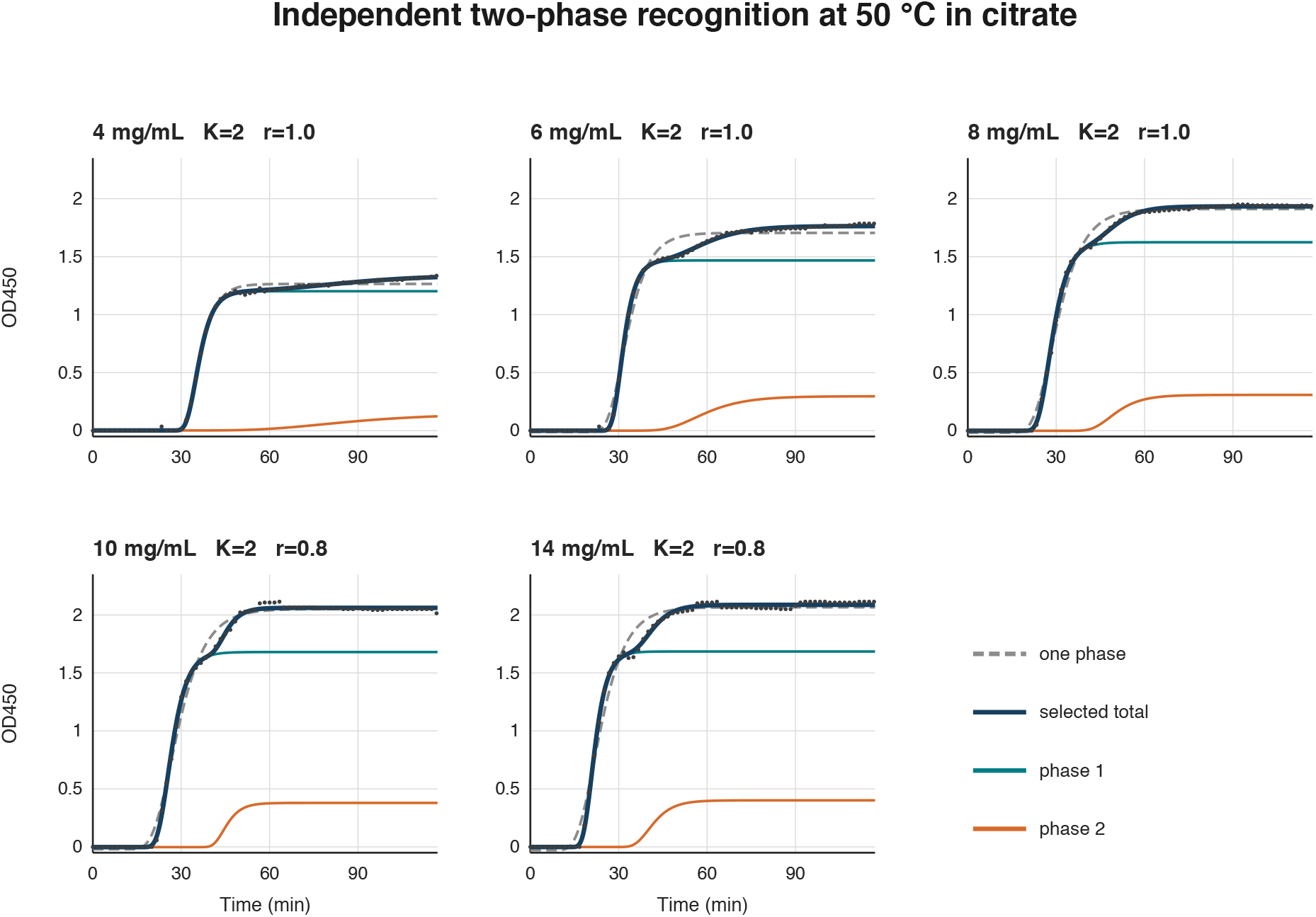
Compact redraw of the 50 °C citrate series. Points are digitized data; grey dashed curves are one-phase fits; navy curves are blind-selected totals; teal and orange curves are blind phase contributions. The 12 mg/mL record is excluded.

**Figure 5:**
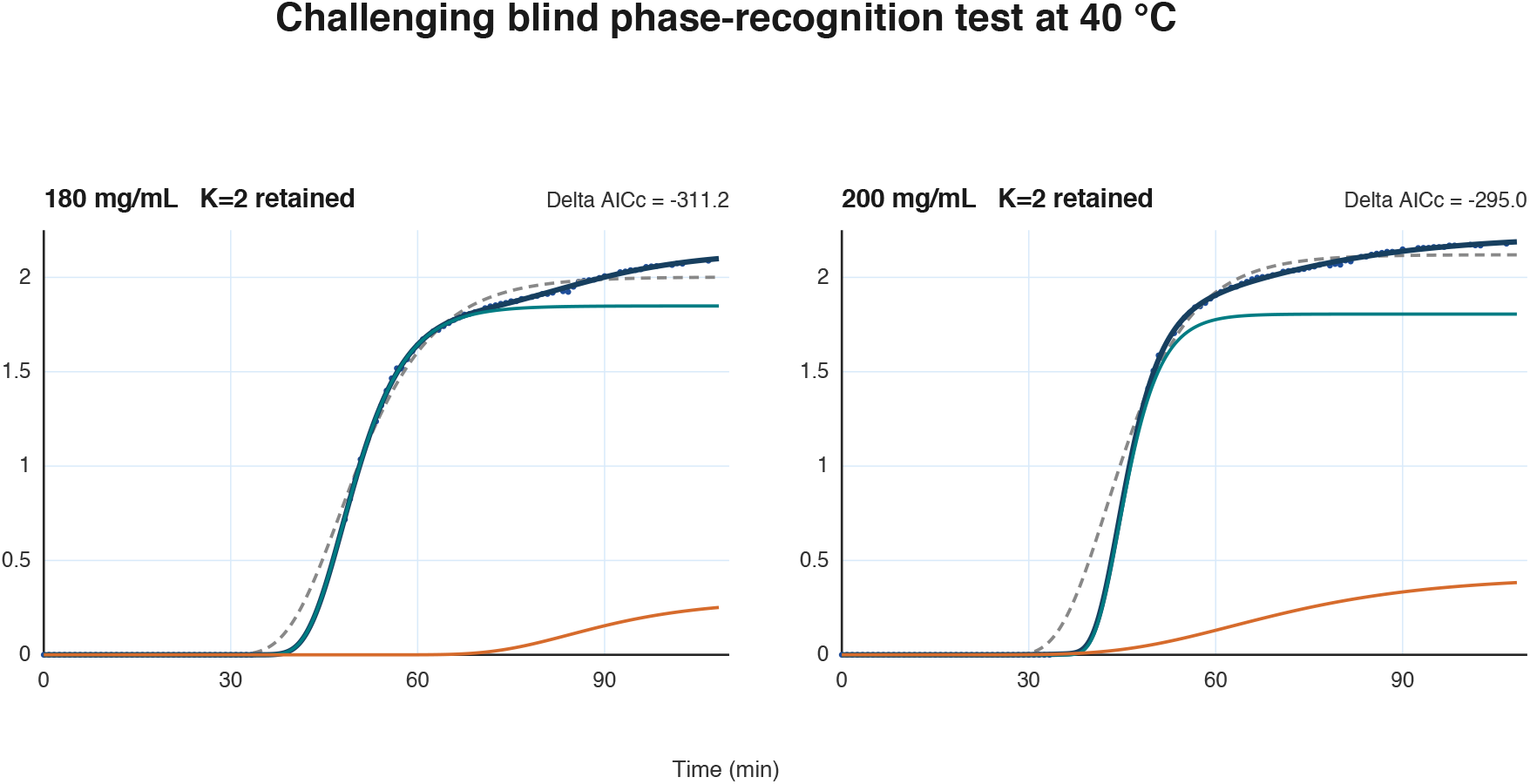
Challenging 40 °C high-concentration test. Points are reconstructed by digitizing the instrument-output plot at its displayed 50-s cadence. Grey dashed curves are one-phase fits; navy curves are blind-selected totals; teal and orange curves are the selected component contributions. The 180 mg/mL trace includes the documented correction of a visible instrument offset at approximately 5350 s. Both illustrative traces retain two phases. Component curves overlap during the leading rise, so the navy total is the relevant representation of fit quality.

#### 3.3.2 Observed Vmax and blind-fit phase rates are distinct estimators

Table 3 compares the reported moving-window Vmax to the blind phase rates. Because moving-window Vmax and an instantaneous Gompertz tangent are different optical estimators, agreement is not expected to be exact; the comparison is descriptive and was not used to select the model. As in the 53 °C series, the same optical comparison is shown on a phase-1-equivalent *µ*M*/*min axis. Blind phase 2 remains optically smaller than phase 1, but its conditional mass-equivalent rate becomes comparable at the higher concentrations and exceeds phase 1 at 10 mg/mL.

**Table 3:**
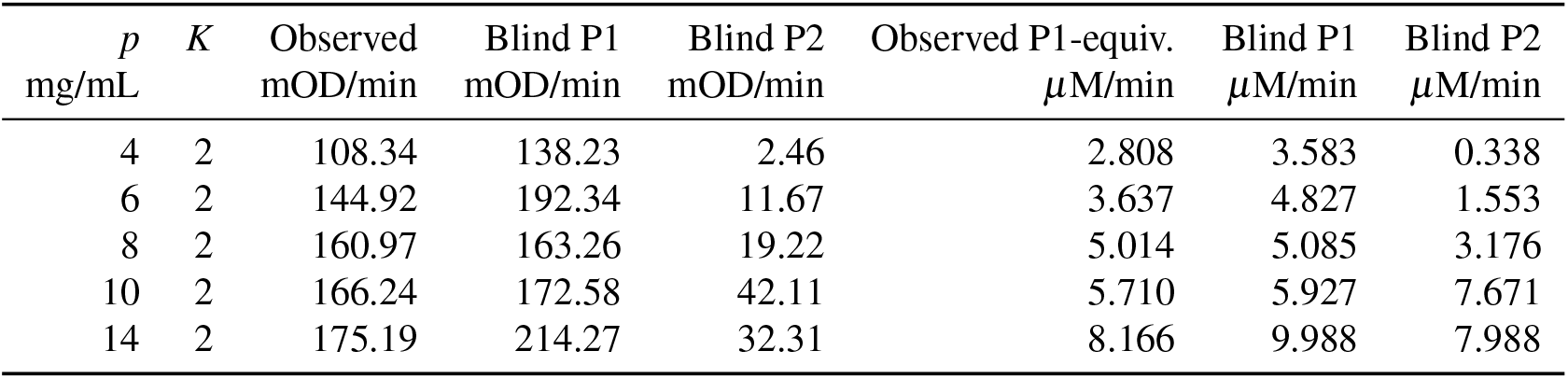
50 °C citrate series: observed versus blind-fit rates. “Observed” is the printed moving-window Vmax. Observed phase-1-equivalent mass rates use same-trace blind phase-1 scaling; all *µ*M*/*min values assume *f*_agg_ = 0.22.

**Table 4:**
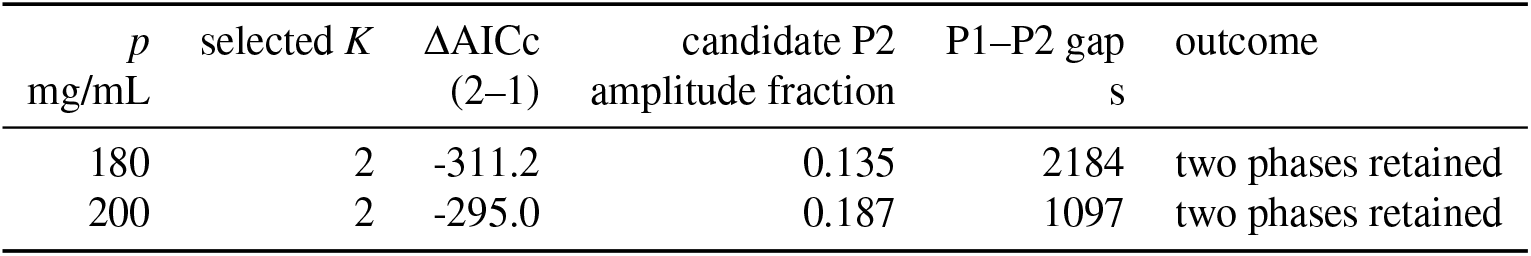
40 °C high-concentration challenge: blind-selection outcomes from digitized instrument-output traces.

### 3.4 Challenging 40 °C high-concentration test

#### 3.4.1 Phase recognition remains informative when transitions are visually obscured

The 40 °C high-concentration traces provide a demanding test because the dominant rise is broad and any late increment is not visually obvious in the original display. These records were analyzed independently, using the digitized instrument-output traces only; no printed Vmax, mass conversion, or result from either primary series entered selection. The hierarchy retained *K* = 2 for both the 180 and 200 mg/mL illustrative traces. The leading shoulder is not fitted perfectly by a single isolated component because both Gompertz terms contribute during the visible rise. Their overlap makes the individual onset parameters correlated, especially after digitization; the selected total, rather than either component alone, is the fitted observable. Thus the phase count is supported, whereas physical interpretation of a component’s earliest tail should remain cautious.

## 4 Discussion

The present paper should be distinguished from several established results. Krishnan and Raibekas already demonstrated a well-defined biphasic IL-1ra aggregation trace and linked its two transitions to Type I and Type II aggregate populations [5]. Other aggregation studies have fitted two sigmoidal terms to resolve sequential populations, including two Boltzmann growth terms for oligomer and fibril formation in immunoglobulin light-chain aggregation [7]; double-sigmoidal behavior has also been documented in insulin aggregation [8]. More generally, multi-sigmoidal Gompertz functions and forward selection of model order have been developed as mathematical growth-curve models [9]. These precedents mean that neither the existence of multiphasic aggregation, nor the use of more than one sigmoidal term, nor information-criterion model selection is claimed here as new.

The distinction of this work is operational. A blind, condition-specific hierarchy is used that starts from one Gompertz component, tests an added component, and continues to a third only when the preceding model is supported by prespecified improvement, amplitude, and temporal-separation gates. The phase count is selected from the turbidity trace alone. Observed rate markers and the Type-I/Type-II response factor are withheld until after selection. The observed-versus-blind comparison is then reported before and after conversion, with the latter explicitly labeled phase-1-equivalent rather than direct mass measurement. In this form, the method is a reproducible phase-recognition workflow for ordinary turbidity data, not a new microscopic aggregation mechanism or a new universal kinetic law.

The principal result is blind recognition of a multiphase optical process at the early concentration of 5 mg/mL in the 53 °C phosphate series. This conclusion uses only the traces and prespecified structural gates. The withheld source rates enter afterward and show that the selected initial component tracks the source optical marker closely. The framework therefore answers a defined practical question: when does one whole-trace turbidity rate stop representing the initial resolved event?

The 40 °C challenge qualifies the scope of this claim. At high concentration and lower temperature, the data display is compatible with a dominant smooth transition and visually subtle late behavior. Nevertheless, the same trace-only hierarchy retains a second component in both illustrative records. Because these values were reconstructed from the instrument-output graphic, this is a challenging demonstration of recognition behavior rather than an independent raw-data validation or a basis for quantitative rate comparison. The small leading-shoulder mismatch is consistent with overlap of the two empirical components and finite raster resolution; it limits mechanistic interpretation of component onset, not the trace-only phase-selection result.

The published Type-I/Type-II response ratio changes the interpretation, but not the recognition, of the selected phases. The late phase can have a modest optical rate while carrying a substantial conditional mass-equivalent rate. This conversion should not be mistaken for direct same-condition mass measurement: the 5.304 response ratio and 22% endpoint aggregate fraction were measured in a different experiment [5]. Every reported *µ*M*/*min value scales with the actual endpoint aggregate fraction under its own condition.

The 53 °C phosphate and 50 °C citrate results should not be combined into a universal concentration-rate relation. They differ in the sole formulation-buffer component (10 mM phosphate versus 10 mM citrate), temperature, and concentration range. Prior IL-1ra work shows that buffer anion can materially change aggregation suppression [6]; it does not provide a phase-specific multiplier that would normalize these two series. The proper next test is matched endpoint mass measurement and matched buffer/temperature experiments, followed by the same blind selection procedure.

## 5 Conclusion

Blind hierarchical selection identifies the first resolved multiphase IL-1ra turbidity process at 5 mg/mL in the 53 °C phosphate series and selects two phases across the retained 50 °C citrate range. In a separate, high-concentration 40 °C challenge, it retains two phases in both illustrative digitized instrument-output traces. Within the two primary series, observed optical markers and blind phase rates are presented side by side in tables before and after a clearly conditional mass-equivalent conversion. The paper’s claim is not a cross-series kinetic law: it is that blind recognition detects early and visually obscured multiphase processes while enforcing structural limits on over-splitting. Because the procedure requires only a turbidity time course and no prior rate or mass calibration, it can be applied directly to other proteins and aggregation assays that produce sigmoidal optical traces, offering a reproducible, low-cost screening step for flagging candidate multiphase behavior before committing to more resource-intensive rate or mass-based characterization.

## Supporting information

Data/code package

## Data and code availability

A companion package accompanies this manuscript as Supplementary Material, containing: the observed-versus-blind rate tables underlying Tables 2 and 3 and their SI counterparts; the fitted Gompertz parameters and derived mass-equivalent quantities for both primary series; the AICc/amplitude/gap values underlying the 40 °C selection summary; the operating-characteristic and perturbation-sensitivity results (Section 2.4); the reconstructed 40 °C digitized trace values with baseline, offset-correction, and bridge-point provenance; and the Python scripts implementing the Gompertz fitting, AICc gating, and validation checks described in Section 2. The raw source image used to digitize the 40 °C instrument-output plot is not included; the digitization script is provided for methodological transparency, and the digitized values themselves are included as data so that all downstream fitting and validation steps remain independently reproducible.

## Companion tool

The blind hierarchical phase-recognition procedure is additionally implemented as a standalone, browser-based tool requiring no installation, archived at Zenodo [11] and available for interactive use at https://www.neurozon.com/tools/blind_phase_recognition.html. Source code is released under the MIT license at https://github.com/neurozi-ops/blind-phase-recognition.

