## Supplemental Info for "Blind Recognition Reveals Early Multiphase IL-1ra Aggregation"

### Supporting Information
### Blind Recognition Reveals Early Multiphase IL-1ra Aggregation

Andrei A. Raibekas*

#### 1 Scope and data separation

This supporting information reports the 53 °C phosphate and 50 °C citrate primary series separately, together with a separate 40 °C high-concentration challenge set (180 and 200 mg/mL). Both primary series use pH 6.5, 140 mM NaCl, and 0.5 mM EDTA. The formulation-buffer component is 10 mM phosphate at 53 °C and 10 mM citrate at 50 °C; temperature and protein-concentration range also differ. The 40 °C set is used for blind phase recognition only, not cross-series rate or mass comparison. No pooled fit, across-series score, or cross-series rate normalization is calculated. The 12 mg/mL citrate trace is omitted throughout.

#### 2 Blind selection rule

For each trace, nested sums of ordered Gompertz components were fitted beginning at one component. A proposed additional component was retained only if: (i) $\Delta$AICc $< -10$; (ii) every component amplitude was greater than 10% of total fitted amplitude; and (iii) every adjacent inflection-time gap exceeded two preceding-component time constants. The 53 °C and 40 °C series considered one or two components. The 50 °C series considered up to three components using ten starts, with at least 80% recurrence required for the selected count. Observed optical rates and the Type-I/Type-II conversion factor were excluded from this selection.

#### 3 Synthetic and sensitivity checks

Four known-ground-truth traces were generated from the ordered Gompertz model: one phase, clearly separated two phases, strongly overlapping two phases, and a two-phase trace with a second amplitude below 10%. Each trace contained 131 points at 50-s spacing. One hundred noisy realizations were fitted using 0.008 OD Gaussian noise and the same selection gates. For a 40 °C-specific operating-characteristic check, 100 one-phase null traces were generated from each trace's fitted $K = 1$ curve and 100 two-phase alternatives from its fitted $K = 2$ curve, preserving the observed time points and adding 0.010 OD Gaussian noise. Separately, 0.010 OD Gaussian perturbations were added pointwise to each digitized 180 and 200 mg/mL trace for 100 complete refits. For 180 mg/mL, the post-offset correction was also drawn from $0.0282 \pm 0.0050$ OD, with separate runs retaining or omitting the linear bridge point. These checks are sensitivity analyses within the fitted Gompertz model family, not estimates of instrument error or independent biological validation.

*Neurozon LLC, 1884 Eastman Avenue, Ventura, CA 93003.

Table 1: Synthetic and digitization-perturbation checks.

| Case | True $K$ | $K = 1$ (%) | $K = 2$ (%) | Median ΔAICc |
|---|---|---|---|---|
| One phase | 1 | 100 | 0 | 4.9 |
| Two phase, separated | 2 | 0 | 100 | -699.9 |
| Two phase, overlapping | 2 | 96 | 4 | -41.9 |
| Two phase, sub-threshold | 2 | 95 | 5 | -351.7 |
| Digitized 180 mg/mL | – | 0 | 100 | -221.1 |
| Digitized 200 mg/mL | – | 0 | 100 | -187.4 |

##### 3.1 40 °C-specific robustness check

Table 2: 40 °C-specific matched simulation and correction-sensitivity checks. Each percentage is based on 100 complete refits. The matched null and alternative traces used the observed time points and 0.010 OD Gaussian noise. For 180 mg/mL, the offset-sensitivity runs varied the correction around 0.0282 OD and either retained or omitted the bridge point.

| Case | 180 $K = 2$ (%) | 180 median ΔAICc | 200 $K = 2$ (%) | 200 median ΔAICc |
|---|---|---|---|---|
| Matched one-phase null | 0 | 4.9 | 0 | 6.6 |
| Matched two-phase alternative | 100 | -255.5 | 99 | -217.4 |
| Digitization perturbation | 100 | -221.4 | 100 | -187.8 |
| 180 offset + bridge sensitivity | 100 | -219.4 | – | – |
| 180 offset + bridge omitted | 100 | -218.5 | – | – |

Table 3: Blind phase-selection summary for the 53 °C phosphate series.

| $p$ mg/mL | selected $K$ | ΔAICc for selected split | source marker mOD/min | outcome |
|---|---|---|---|---|
| 1 | 1 | split rejected | 104.6 | temporal separation failed |
| 2 | 1 | split rejected | 237.4 | information support failed |
| 5 | 2 | -173.2 | 408.0 | two phases retained |
| 10 | 2 | -154.7 | 552.5 | two phases retained |

#### 4 40 °C high-concentration challenge set

The available 40 °C record is a high-resolution SigmaPlot-style graphic derived from the instrument output, rather than a native numerical export. Marker coordinates for the retained 180 and 200 mg/mL traces were digitized at the displayed 50-s cadence. The 180 mg/mL trace contains a visible positive offset at approximately 5350 s; a 0.028 OD post-offset correction and a single linear bridge across the discontinuity were applied before fitting. The accompanying CSV identifies each baseline, digitized, corrected, and bridge value. This set is therefore a challenging test of phase recognition, not an independent raw-data validation. The individual empirical components overlap over the leading rise, so their earliest onset parameters are correlated; the selected total is the fitted observable. As a fit-quality diagnostic, the selected two-phase total reduced RMSE from 0.0368 to 0.0073 OD at 180 mg/mL and from 0.0293 to 0.0062 OD at 200 mg/mL relative to the one-phase fit. These residual reductions do not by themselves establish two biological mechanisms.

Table 4: Blind phase-selection summary for the 40 °C high-concentration challenge.

| $p$ mg/mL | selected $K$ | ΔAICc (2–1) | candidate P2 amplitude fraction | P1–P2 gap s | outcome |
|---|---|---|---|---|---|
| 180 | 2 | -311.2 | 0.135 | 2184 | two phases retained |
| 200 | 2 | -295.0 | 0.187 | 1097 | two phases retained |

Table 5: Blind phase-selection summary for the 50 °C citrate series. Printed Vmax values were not used in selection.

| $p$ mg/mL | selected $K$ | recurrence starts/10 | printed Vmax mOD/min | outcome |
|---|---|---|---|---|
| 4 | 2 | 10 | 108.34 | third phase unresolved |
| 6 | 2 | 10 | 144.92 | third phase too small |
| 8 | 2 | 10 | 160.97 | third phase too small |
| 10 | 2 | 8 | 166.24 | third phase too small |
| 14 | 2 | 8 | 175.19 | third phase too small |

### 5 Observed-versus-blind rate comparisons

The observed optical comparator is a withheld source marker in the 53 °C series and a printed moving-window Vmax in the 50 °C series. It is not a phase-specific observation and was not used by the blind procedure. Blind P1 and P2 are instantaneous Gompertz peak rates. The observed phase-1-equivalent $\mu\text{M/min}$ value is obtained by multiplying the observed optical rate by the same-trace blind-P1 mass/optical ratio. It preserves the observed-versus-fit comparison after conversion but is not an independently measured mass rate.

Table 6: 53 °C phosphate series: observed and blind-fit rates before and after conversion. All mass-equivalent rates assume $f_{\text{agg}} = 0.22$.

| $p$ mg/mL | $K$ | observed mOD/min | blind P1 mOD/min | blind P2 mOD/min | observed P1-equiv. $\mu$M/min | blind P1 $\mu$M/min | blind P2 $\mu$M/min |
|---|---|---|---|---|---|---|---|
| 1 | 1 | 104.60 | 103.85 | – | 1.531 | 1.520 | – |
| 2 | 1 | 237.40 | 234.24 | – | 4.434 | 4.375 | – |
| 5 | 2 | 408.00 | 411.50 | 40.90 | 6.806 | 6.864 | 3.619 |
| 10 | 2 | 552.50 | 563.36 | 97.59 | 16.336 | 16.657 | 15.305 |

### 6 Mass-equivalent calculation

The Krishnan–Raibekas study [1] reports a 5.304-fold $\text{OD}_{450}$ response-per-mass ratio for Type I relative to Type II IL-1ra aggregate material. Blind phase 1 is assigned to the higher-response population and blind phase 2 to the lower-response population only for post-selection conversion. For a two-phase trace,

$$F_1 = \frac{a_1/5.304}{a_1/5.304 + a_2}, \qquad F_2 = \frac{a_2}{a_1/5.304 + a_2}. \tag{1}$$

With molecular weight 17.3 kDa, $C_{\text{tot}} = 57.803p$ $\mu$M for $p$ in mg/mL. The conditional peak mass-equivalent rate is

$$v_{j,\max} = \frac{0.22 C_{\text{tot}} F_j}{e b_j} \quad (\mu\text{M/min}). \tag{2}$$

The sample volume is 0.2 mL, so the amount rate is $0.2 v_{j,\max}$ nmol/min. If the actual endpoint aggregate fraction is $f$, all listed mass-equivalent rates scale by $f/0.22$. The response factor and endpoint fraction are published external quantities, not measurements in either present kinetic series.

Table 7: 50 °C citrate series: observed and blind-fit rates before and after conversion. All mass-equivalent rates assume $f_{\text{agg}} = 0.22$.

| $p$ mg/mL | $K$ | observed mOD/min | blind P1 mOD/min | blind P2 mOD/min | observed P1-equiv. $\mu$M/min | blind P1 $\mu$M/min | blind P2 $\mu$M/min |
|---|---|---|---|---|---|---|---|
| 4 | 2 | 108.34 | 138.23 | 2.46 | 2.808 | 3.583 | 0.338 |
| 6 | 2 | 144.92 | 192.34 | 11.67 | 3.637 | 4.827 | 1.553 |
| 8 | 2 | 160.97 | 163.26 | 19.22 | 5.014 | 5.085 | 3.176 |
| 10 | 2 | 166.24 | 172.58 | 42.11 | 5.710 | 5.927 | 7.671 |
| 14 | 2 | 175.19 | 214.27 | 32.31 | 8.166 | 9.988 | 7.988 |
