## Supplementary material for "Blind Recognition Reveals Early Multiphase IL-1ra Aggregation": Data/code package: analyze_40c_challenge.py

```
"""Blind two-phase challenge analysis for digitized high-concentration 40 °C traces."""

from pathlib import Path
import csv
import sys

import numpy as np
from reportlab.lib.colors import Color, HexColor
from reportlab.pdfgen import canvas
import pypdfium2 as pdfium

HERE = Path(__file__).resolve().parent
ROOT = HERE.parents[1]
sys.path.insert(0, str(HERE))
from analyze_slide4 import fit_model, gompertz  # reuse the prespecified fitter

DATA = ROOT / "outputs" / "IL1ra_40C_digitized_traces.csv"
OUT_CSV = ROOT / "outputs" / "hierarchical_selection_summary_40C.csv"
OUT_PDF = ROOT / "outputs" / "figs_v2" / "fig5_40C_challenging_test.pdf"
OUT_PNG = ROOT / "outputs" / "figs_v2" / "fig5_40C_challenging_test.png"
CONCENTRATIONS = (180, 200)


def load_data():
    data = {c: [[], []] for c in CONCENTRATIONS}
    with DATA.open() as f:
        for row in csv.DictReader(f):
            c = int(row["concentration_mg_ml"])
            if c in data:
                data[c][0].append(float(row["time_s"]))
                data[c][1].append(float(row["od_digitized_corrected"]))
    return {c: (np.asarray(v[0]), np.asarray(v[1])) for c, v in data.items()}


def assess(t, y):
    one = fit_model(t, y, 1)
    two = fit_model(t, y, 2)
    p = two[2]
    a1, a2 = float(p[1]), float(p[4])
    amp_total = a1 + a2
    amp2_fraction = a2 / amp_total
    gap = float(p[5] - p[2])
    width1 = float(p[3])
    delta = float(two[1] - one[1])
    selected = delta < -10 and min(a1, a2) / amp_total >= 0.10 and gap > 2 * width1
    return {
        "one": one, "two": two, "delta": delta, "amp2_fraction": amp2_fraction,
        "gap": gap, "width1": width1, "selected": selected,
    }


def _line(c, xy, xs, ys, color, width=1.0, dashed=False):
    c.setStrokeColor(HexColor(color)); c.setLineWidth(width)
    c.setDash(3, 2) if dashed else c.setDash()
    path = c.beginPath()
    for i, (xv, yv) in enumerate(zip(xs, ys)):
        xx, yy = xy(xv, yv)
        path.moveTo(xx, yy) if i == 0 else path.lineTo(xx, yy)
    c.drawPath(path, stroke=1, fill=0); c.setDash()


def _panel(c, x, y, title):
    w, h = 213, 132
    for xv in (0, 30, 60, 90):
        xx = x + xv / 110 * w
        c.setStrokeColor(HexColor("#d9eafa")); c.setLineWidth(0.4); c.line(xx, y, xx, y + h)
        c.setFillColor(Color(.2, .2, .2)); c.setFont("Helvetica", 6.5); c.drawCentredString(xx, y - 10, str(xv))
    for yv in (0, .5, 1, 1.5, 2):
        yy = y + yv / 2.25 * h
        c.setStrokeColor(HexColor("#d9eafa")); c.setLineWidth(0.4); c.line(x, yy, x + w, yy)
        c.setFillColor(Color(.2, .2, .2)); c.setFont("Helvetica", 6.5); c.drawRightString(x - 4, yy - 2, f"{yv:g}")
    c.setStrokeColor(Color(.15, .15, .15)); c.setLineWidth(.7); c.line(x, y, x + w, y); c.line(x, y, x, y + h)
    c.setFillColor(Color(.1, .1, .1)); c.setFont("Helvetica-Bold", 8); c.drawString(x, y + h + 7, title)
    return lambda xv, yv: (x + xv / 110 * w, y + yv / 2.25 * h)


def plot(data, results):
    OUT_PDF.parent.mkdir(parents=True, exist_ok=True)
    width, height = 540, 270
    c = canvas.Canvas(str(OUT_PDF), pagesize=(width, height))
    c.setFillColor(Color(.1, .1, .1)); c.setFont("Helvetica-Bold", 12)
    c.drawCentredString(width / 2, height - 20, "Challenging blind phase-recognition test at 40 °C")
    positions = [(58, 56), (306, 56)]
    for (x, y0), conc in zip(positions, CONCENTRATIONS):
        t, y = data[conc]; r = results[conc]
        label = "K=2 retained" if r["selected"] else "K=1 retained"
        xy = _panel(c, x, y0, f"{conc} mg/mL   {label}")
        c.setFillColor(Color(.2, .2, .2)); c.setFont("Helvetica", 6.5)
        # Put the statistic on the title line, outside the plotting frame, so
        # it cannot cover the late trace or the phase-2 shoulder.
        c.drawRightString(x + 213, y0 + 139, f"Delta AICc = {r['delta']:.1f}")
        c.setFillColor(HexColor("#1f4e9a"))
        for tx, oy in zip(t / 60, y):
            px, py = xy(tx, oy); c.circle(px, py, .9, stroke=0, fill=1)
        dense = np.linspace(0, 6500, 450)
        _line(c, xy, dense / 60, gompertz(dense, r["one"][2], 1), "#888888", 1.0, True)
        if r["selected"]:
            p = r["two"][2]; base = p[0]
            _line(c, xy, dense / 60, gompertz(dense, p, 2), "#173f5f", 1.65)
            _line(c, xy, dense / 60, base + p[1] * np.exp(-np.exp(-(dense-p[2])/p[3])), "#007c83", 1.0)
            _line(c, xy, dense / 60, p[4] * np.exp(-np.exp(-(dense-p[5])/p[6])), "#d66b2c", 1.0)
    c.setFillColor(Color(.2, .2, .2)); c.setFont("Helvetica", 7)
    c.drawCentredString(width / 2, 18, "Time (min)")
    c.save()
    page = pdfium.PdfDocument(str(OUT_PDF))[0]
    page.render(scale=2.2).to_pil().save(OUT_PNG)


def main():
    data = load_data()
    results = {c: assess(*data[c]) for c in CONCENTRATIONS}
    with OUT_CSV.open("w", newline="") as f:
        writer = csv.writer(f)
        writer.writerow(["concentration_mg_ml", "n_digitized", "one_aicc", "two_aicc",
                         "delta_aicc_2_minus_1", "phase2_amplitude_fraction",
                         "inflection_gap_s", "phase1_width_s", "selected_phase_count"])
        for c in CONCENTRATIONS:
            r = results[c]
            writer.writerow([c, len(data[c][0]), f"{r['one'][1]:.3f}", f"{r['two'][1]:.3f}",
                             f"{r['delta']:.3f}", f"{r['amp2_fraction']:.3f}",
                             f"{r['gap']:.1f}", f"{r['width1']:.1f}", 2 if r["selected"] else 1])
    plot(data, results)
    for c in CONCENTRATIONS:
        r = results[c]
        print(c, "K=", 2 if r["selected"] else 1, "dAICc=", round(r["delta"], 2),
              "amp2=", round(r["amp2_fraction"], 3), "gap=", round(r["gap"], 1),
              "width1=", round(r["width1"], 1))


if __name__ == "__main__":
    main()
```
