## Supplementary material for "Blind Recognition Reveals Early Multiphase IL-1ra Aggregation": Data/code package: analyze_slide4.py

```
"""Exploratory blind phase-count test on the rasterized 40 C slide-4 chart."""

from pathlib import Path

import numpy as np
from PIL import Image


ROOT = Path(__file__).resolve().parent
IMAGE = ROOT / "source" / "slide-4.png"
OUT = ROOT / "slide4_digitized_rates.csv"

# Original slide-4 chart coordinates, read from the rendered chart.
X_LEFT, X_RIGHT = 523.0, 1148.0
Y_TOP, Y_BOTTOM = 222.0, 520.0
T_MAX, OD_MAX = 6500.0, 2.2
# Keep all plotted curves while digitizing because their vertical ordering
# identifies their concentration. Report only the four retained challenge
# traces after that extraction.
ALL_CONCENTRATIONS = np.array([200.0, 180.0, 160.0, 140.0, 120.0, 100.0])
CONCENTRATIONS = np.array([200.0, 180.0, 160.0, 120.0])
PRINTED_VMAX = {200.0: 121.50, 180.0: 100.79, 160.0: 90.479,
                140.0: 69.959, 120.0: 47.001, 100.0: 25.889}


def pixel_to_time(x):
    return (x - X_LEFT) / (X_RIGHT - X_LEFT) * T_MAX


def pixel_to_od(y):
    return (Y_BOTTOM - y) / (Y_BOTTOM - Y_TOP) * OD_MAX


def blue_y_clusters(image, x, half_width=2, cluster_gap=8):
    """Return centers of dark-blue trace pixels in a narrow vertical strip."""
    ys = []
    for xx in range(int(x - half_width), int(x + half_width + 1)):
        p = image[:, xx, :]
        mask = (
            (p[:, 0] < 100) & (p[:, 1] < 100) & (p[:, 2] > 150)
            & (p[:, 2] > p[:, 0] + 60) & (p[:, 2] > p[:, 1] + 60)
        )
        ys.extend(np.where(mask)[0].tolist())
    ys = sorted(ys)
    clusters = []
    for y in ys:
        if not clusters or y - clusters[-1][-1] > cluster_gap:
            clusters.append([y])
        else:
            clusters[-1].append(y)
    centers = [float(np.mean(c)) for c in clusters]
    return [y for y in centers if Y_TOP - 5 < y < Y_BOTTOM + 5]


def digitize_traces(x_start=800, x_end=1121, cluster_gap=8):
    image = np.asarray(Image.open(IMAGE).convert("RGB"))
    x_values = np.arange(x_start, x_end, 3)
    observations = {float(c): [] for c in ALL_CONCENTRATIONS}

    # At sufficiently separated times the six non-crossing curves are ordered
    # top-to-bottom by concentration. Keep only strips with six trace centers,
    # avoiding marker artifacts and unresolved early overlap.
    for x in x_values:
        centers = blue_y_clusters(image, x, cluster_gap=cluster_gap)
        if len(centers) != 6:
            continue
        for c, y in zip(ALL_CONCENTRATIONS, centers):
            observations[float(c)].append((pixel_to_time(x), pixel_to_od(y)))

    # The common pre-transition baseline is visually at OD=0. It is included
    # only to anchor the digitized trace; no source rate enters fitting.
    for c in ALL_CONCENTRATIONS:
        baseline = [(float(t), 0.0) for t in np.arange(0, 2001, 100)]
        observations[float(c)] = baseline + observations[float(c)]
    return {float(c): observations[float(c)] for c in CONCENTRATIONS}


def gompertz(t, params, k):
    baseline = params[0]
    values = np.full_like(t, baseline, dtype=float)
    for j in range(k):
        amp, t0, width = params[1 + 3 * j: 4 + 3 * j]
        values += amp * np.exp(-np.exp(-(t - t0) / width))
    return values


def fit_model(t, y, k):
    max_y = max(float(np.max(y)), 0.1)
    starts = []
    if k == 1:
        for t0 in (2400, 2700, 3000, 3300):
            starts.append([0.0, max_y, t0, 500.0])
    else:
        for t01, t02 in ((2450, 3350), (2550, 3600), (2750, 3900), (3000, 4200)):
            starts.append([0.0, max_y * 0.65, t01, 350.0, max_y * 0.35, t02, 700.0])

    if k == 1:
        lower = [-0.08, 0.01, 1500.0, 80.0]
        upper = [0.08, 3.0, 5500.0, 4000.0]
    else:
        lower = [-0.08, 0.01, 1500.0, 80.0, 0.01, 1500.0, 80.0]
        upper = [0.08, 3.0, 5500.0, 4000.0, 3.0, 6000.0, 4000.0]

    def objective(params):
        params = np.asarray(params)
        if np.any(params < np.asarray(lower)) or np.any(params > np.asarray(upper)):
            return 1e9 + float(np.sum(np.maximum(np.asarray(lower) - params, 0) ** 2))
        if k == 2 and params[2] > params[5]:
            return 1e8 + (params[2] - params[5]) ** 2
        residual = gompertz(t, params, k) - y
        return float(np.sum(residual ** 2))

    def nelder_mead(start, scales, max_iter=1800):
        start = np.asarray(start, dtype=float)
        simplex = [start]
        for i, scale in enumerate(scales):
            trial = start.copy()
            trial[i] += scale
            simplex.append(trial)
        values = [objective(point) for point in simplex]
        for _ in range(max_iter):
            order = np.argsort(values)
            simplex = [simplex[i] for i in order]
            values = [values[i] for i in order]
            if np.max(np.abs(np.asarray(values) - values[0])) < 1e-8:
                break
            centroid = np.mean(simplex[:-1], axis=0)
            worst = simplex[-1]
            reflected = centroid + (centroid - worst)
            reflected_value = objective(reflected)
            if values[0] <= reflected_value < values[-2]:
                simplex[-1], values[-1] = reflected, reflected_value
                continue
            if reflected_value < values[0]:
                expanded = centroid + 2.0 * (reflected - centroid)
                expanded_value = objective(expanded)
                simplex[-1], values[-1] = (expanded, expanded_value) if expanded_value < reflected_value else (reflected, reflected_value)
                continue
            contracted = centroid + 0.5 * (worst - centroid)
            contracted_value = objective(contracted)
            if contracted_value < values[-1]:
                simplex[-1], values[-1] = contracted, contracted_value
                continue
            best_point = simplex[0]
            simplex = [best_point + 0.5 * (point - best_point) for point in simplex]
            values = [objective(point) for point in simplex]
        best_index = int(np.argmin(values))
        return float(values[best_index]), simplex[best_index]

    best = None
    for start in starts:
        scales = [0.02, 0.15, 120.0, 80.0] if k == 1 else [0.02, 0.15, 120.0, 80.0, 0.15, 120.0, 160.0]
        rss, params = nelder_mead(start, scales)
        if k == 2 and params[2] > params[5]:
            params = np.array([params[0], params[4], params[5], params[6], params[1], params[2], params[3]])
        if best is None or rss < best[0]:
            best = (rss, params)

    rss, params = best
    n = len(y)
    p = len(params)
    aic = n * np.log(max(rss / n, 1e-12)) + 2 * p
    aicc = aic + (2 * p * (p + 1)) / max(n - p - 1, 1)
    return rss, aicc, params


def main():
    observations = digitize_traces()
    rows = []
    for c in CONCENTRATIONS:
        data = np.asarray(observations[float(c)])
        t, y = data[:, 0], data[:, 1]
        one = fit_model(t, y, 1)
        two = fit_model(t, y, 2)
        total_amp = float(np.sum(two[2][1::3]))
        amp1 = float(two[2][1] / total_amp)
        amp2 = float(two[2][4] / total_amp)
        t_gap = float(two[2][5] - two[2][2])
        rows.append((c, len(t), one[0], two[0], one[1], two[1], two[1] - one[1],
                     PRINTED_VMAX[float(c)], amp1, amp2, t_gap, float(two[2][3])))

    header = "concentration_mg_ml,n_digitized,one_rss,two_rss,one_aicc,two_aicc,delta_aicc_2_minus_1,printed_vmax_mOD_min,two_amp1_fraction,two_amp2_fraction,two_t0_gap_sec,two_phase1_width_sec"
    lines = [header]
    for row in rows:
        lines.append(",".join(f"{v:.8g}" if isinstance(v, float) else str(v) for v in row))
    OUT.write_text("\n".join(lines) + "\n")
    print(header)
    print("\n".join(lines[1:]))


if __name__ == "__main__":
    main()
```
