## Supplementary material for "Blind Recognition Reveals Early Multiphase IL-1ra Aggregation": Data/code package: digitize_40c_highres.py

```
"""Reconstruct the plotted 40 C OD traces from the highest-resolution raster.

This is deliberately a digitization workflow, not a claim to recover the
instrument export.  The 180 mg/mL curve contains a visible instrument offset
near 5,300 s.  The common post-offset displacement is removed so its series is
continuous with the immediately preceding trace segment.
"""

from pathlib import Path
import csv

import numpy as np
from PIL import Image


SOURCE = Path("/Users/andreir/Desktop/Aggregation/IL-1ra 40C traces.png")
OUT = Path(__file__).resolve().parents[2] / "outputs" / "IL1ra_40C_digitized_traces.csv"

# Calibrated against the labelled plot frame: 0--6500 s and 0--2.2 OD.
X_LEFT, X_RIGHT = 398.0, 2114.0
Y_TOP, Y_BOTTOM = 56.0, 875.0
T_MAX, OD_MAX = 6500.0, 2.2
ALL_CONCENTRATIONS = (200, 180, 160, 140, 120, 100)
RETAINED = (200, 180, 160, 120)
CADENCE_S = 50


def x_from_time(t):
    return X_LEFT + t / T_MAX * (X_RIGHT - X_LEFT)


def od_from_y(y):
    return (Y_BOTTOM - y) / (Y_BOTTOM - Y_TOP) * OD_MAX


def blue_centers(image, x, half_width=3, gap=12):
    """Find vertically separated blue traces near a plotted marker column."""
    ys = []
    for xx in range(round(x - half_width), round(x + half_width) + 1):
        pixel = image[:, xx, :]
        mask = (
            (pixel[:, 0] < 80) & (pixel[:, 1] < 120) & (pixel[:, 2] > 130)
            & (pixel[:, 2] > pixel[:, 0] + 55)
            & (pixel[:, 2] > pixel[:, 1] + 45)
        )
        ys.extend(np.flatnonzero(mask).tolist())
    ys.sort()
    clusters = []
    for y in ys:
        if not clusters or y - clusters[-1][-1] > gap:
            clusters.append([y])
        else:
            clusters[-1].append(y)
    return [np.mean(c) for c in clusters if Y_TOP - 4 < np.mean(c) < Y_BOTTOM + 4]


def extract():
    image = np.asarray(Image.open(SOURCE).convert("RGB"))
    series = {c: [] for c in ALL_CONCENTRATIONS}
    # The marker cadence is 50 s (ten observed points per 500-s major grid).
    # Baseline points cannot
    # be separated by concentration before the traces diverge, so those shared
    # baseline observations are recorded as OD=0.
    for t in range(0, 2001, CADENCE_S):
        for c in ALL_CONCENTRATIONS:
            series[c].append((float(t), 0.0, "shared_baseline"))
    for t in range(2050, 6501, CADENCE_S):
        centers = blue_centers(image, x_from_time(t))
        if len(centers) != 6:
            continue
        for c, y in zip(ALL_CONCENTRATIONS, centers):
            series[c].append((float(t), float(od_from_y(y)), "digitized"))
    return series


def correct_180_offset(points):
    """Remove the visible +OD step beginning at approximately 5,340 s.

    The magnitude is determined from local straight-line extrapolations on
    either side of the discontinuity.  Applying the common offset to the
    post-step values preserves the local shape while removing the artefact.
    """
    numeric = [(t, od) for t, od, _ in points if 5100 <= t <= 5300 or 5350 <= t <= 5600]
    pre = np.array([(t, od) for t, od in numeric if t <= 5300])
    post = np.array([(t, od) for t, od in numeric if t >= 5350])
    if len(pre) < 2 or len(post) < 2:
        return points, 0.0
    pre_line = np.polyfit(pre[:, 0], pre[:, 1], 1)
    post_line = np.polyfit(post[:, 0], post[:, 1], 1)
    boundary = 5325.0
    jump = float(np.polyval(post_line, boundary) - np.polyval(pre_line, boundary))
    corrected = []
    for t, od, source in points:
        if t >= 5350 and source == "digitized":
            corrected.append((t, od - jump, "digitized; 180 mg/mL offset corrected"))
        else:
            corrected.append((t, od, source))
    # The marker at the discontinuity itself is visually merged with nearby
    # traces.  Insert its value only as the user-specified straight-line bridge,
    # rather than inventing a curved measurement.
    known = {t: (od, source) for t, od, source in corrected}
    if 5350.0 not in known and 5300.0 in known and 5400.0 in known:
        left, right = known[5300.0][0], known[5400.0][0]
        known[5350.0] = ((left + right) / 2, "straight-line bridge across instrument offset")
    corrected = [(t, *known[t]) for t in sorted(known)]
    return corrected, jump


def main():
    data = extract()
    data[180], jump = correct_180_offset(data[180])
    OUT.parent.mkdir(parents=True, exist_ok=True)
    with OUT.open("w", newline="") as f:
        writer = csv.writer(f)
        writer.writerow([
            "concentration_mg_ml", "time_s", "od_digitized_corrected",
            "source_note", "180_offset_removed_od"
        ])
        for c in RETAINED:
            for t, od, note in data[c]:
                writer.writerow([c, f"{t:.0f}", f"{od:.5f}", note,
                                 f"{jump:.5f}" if c == 180 else ""])
    print(f"Wrote {OUT}")
    print(f"180 mg/mL post-5,340 s offset removed: {jump:.4f} OD")
    for c in RETAINED:
        print(c, len(data[c]))


if __name__ == "__main__":
    main()
```
