## Supplementary material for "Blind Recognition Reveals Early Multiphase IL-1ra Aggregation": Data/code package: explore_40c_simulation.py

```
"""Exploratory simulation checks for the digitized 40 C challenge traces.

This script does not modify the manuscript.  It asks three questions:
1. How often would a matched one-phase null be falsely split into two phases?
2. How often would a matched two-phase trace be recovered?
3. How sensitive is the 180 mg/mL result to digitization noise, the bridge point,
   and uncertainty in the documented offset correction?
"""

from __future__ import annotations

import csv
import sys
from pathlib import Path

import numpy as np
from PIL import Image, ImageDraw

HERE = Path(__file__).resolve().parent
ROOT = HERE.parents[1]
FIT_DIR = HERE if (HERE / "analyze_slide4.py").exists() else ROOT / "work" / "slide4_validation"
sys.path.insert(0, str(FIT_DIR))
from analyze_slide4 import fit_model, gompertz  # noqa: E402


DATA = HERE / "IL1ra_40C_digitized_traces.csv"
if not DATA.exists():
    DATA = ROOT / "outputs" / "IL1ra_40C_digitized_traces.csv"
OUT_CSV = HERE / "explore_40c_simulation_summary.csv"
OUT_PNG = HERE / "explore_40c_simulation_summary.png"

CONCENTRATIONS = (180, 200)
N_REPS = 100
SIGMA_OD = 0.010
BASE_SEED = 4082026
OFFSET_NOMINAL = 0.02820


def load_data() -> dict[int, tuple[np.ndarray, np.ndarray]]:
    data = {c: [[], []] for c in CONCENTRATIONS}
    with DATA.open() as f:
        for row in csv.DictReader(f):
            c = int(row["concentration_mg_ml"])
            if c in data:
                data[c][0].append(float(row["time_s"]))
                data[c][1].append(float(row["od_digitized_corrected"]))
    return {c: (np.asarray(v[0]), np.asarray(v[1])) for c, v in data.items()}


def assess(t: np.ndarray, y: np.ndarray) -> dict[str, float | bool]:
    one = fit_model(t, y, 1)
    two = fit_model(t, y, 2)
    p = two[2]
    a1, a2 = float(p[1]), float(p[4])
    total = a1 + a2
    gap = float(p[5] - p[2])
    width1 = float(p[3])
    delta = float(two[1] - one[1])
    selected = delta < -10 and min(a1, a2) / total >= 0.10 and gap > 2 * width1
    return {
        "selected": bool(selected),
        "delta": delta,
        "amp2_fraction": a2 / total,
        "gap": gap,
        "width1": width1,
    }


def summarize(case: str, concentration: int, records: list[dict[str, float | bool]]) -> dict[str, object]:
    delta = np.asarray([float(r["delta"]) for r in records])
    amp2 = np.asarray([float(r["amp2_fraction"]) for r in records])
    gap = np.asarray([float(r["gap"]) for r in records])
    selected = np.asarray([bool(r["selected"]) for r in records])
    return {
        "case": case,
        "concentration_mg_ml": concentration,
        "n": len(records),
        "selected_k2_pct": 100 * float(np.mean(selected)),
        "median_delta_aicc": float(np.median(delta)),
        "p05_delta_aicc": float(np.percentile(delta, 5)),
        "p95_delta_aicc": float(np.percentile(delta, 95)),
        "median_amp2_fraction": float(np.median(amp2)),
        "p05_amp2_fraction": float(np.percentile(amp2, 5)),
        "p95_amp2_fraction": float(np.percentile(amp2, 95)),
        "median_gap_s": float(np.median(gap)),
        "p05_gap_s": float(np.percentile(gap, 5)),
        "p95_gap_s": float(np.percentile(gap, 95)),
    }


def varied_180_trace(t: np.ndarray, y: np.ndarray, rng: np.random.Generator, *,
                     omit_bridge: bool = False) -> tuple[np.ndarray, np.ndarray]:
    """Apply a plausible alternate offset correction and digitization noise."""
    y_var = y.copy()
    # The published reconstructed trace is corrected with OFFSET_NOMINAL.  If
    # the true step were c instead, the corrected post-offset values would be
    # shifted by (nominal - c).  Use a conservative 0.005 OD uncertainty.
    c = float(rng.normal(OFFSET_NOMINAL, 0.005))
    post = t >= 5400
    y_var[post] += OFFSET_NOMINAL - c
    bridge = np.where(t == 5350)[0]
    if len(bridge) and not omit_bridge:
        i = int(bridge[0])
        before = np.where(t == 5300)[0]
        after = np.where(t == 5400)[0]
        if len(before) and len(after):
            y_var[i] = 0.5 * (y_var[int(before[0])] + y_var[int(after[0])])
    if omit_bridge:
        keep = t != 5350
        t = t[keep]
        y_var = y_var[keep]
    y_var += rng.normal(0.0, SIGMA_OD, size=len(y_var))
    return t, y_var


def main() -> None:
    data = load_data()
    rows: list[dict[str, object]] = []
    plot_data: list[tuple[str, int, float]] = []

    for concentration in CONCENTRATIONS:
        t, y = data[concentration]
        one = fit_model(t, y, 1)
        two = fit_model(t, y, 2)
        one_mean = gompertz(t, one[2], 1)
        two_mean = gompertz(t, two[2], 2)
        print(f"{concentration} mg/mL observed: delta AICc = {two[1] - one[1]:.2f}")

        rng = np.random.default_rng(BASE_SEED + concentration)
        cases: list[tuple[str, list[dict[str, float | bool]]]] = []
        null_records = [assess(t, one_mean + rng.normal(0, SIGMA_OD, len(t))) for _ in range(N_REPS)]
        cases.append(("matched one-phase null", null_records))
        alternative_records = [assess(t, two_mean + rng.normal(0, SIGMA_OD, len(t))) for _ in range(N_REPS)]
        cases.append(("matched two-phase alternative", alternative_records))

        perturb_records = [assess(t, y + rng.normal(0, SIGMA_OD, len(t))) for _ in range(N_REPS)]
        cases.append(("digitization perturbation", perturb_records))

        if concentration == 180:
            offset_records = []
            no_bridge_records = []
            for _ in range(N_REPS):
                t_var, y_var = varied_180_trace(t, y, rng, omit_bridge=False)
                offset_records.append(assess(t_var, y_var))
                t_var, y_var = varied_180_trace(t, y, rng, omit_bridge=True)
                no_bridge_records.append(assess(t_var, y_var))
            cases.append(("180 offset + bridge sensitivity", offset_records))
            cases.append(("180 offset + bridge omitted", no_bridge_records))

        for case, records in cases:
            row = summarize(case, concentration, records)
            rows.append(row)
            plot_data.append((case, concentration, float(row["selected_k2_pct"])))

    fieldnames = list(rows[0].keys())
    with OUT_CSV.open("w", newline="") as f:
        writer = csv.DictWriter(f, fieldnames=fieldnames)
        writer.writeheader()
        writer.writerows(rows)

    # Compact selection-probability plot using only Pillow, which is available
    # in the bundled runtime.  This is diagnostic output, not a manuscript
    # figure.
    labels = ["one-phase null", "two-phase alternative", "digitization perturbation",
              "offset + bridge", "offset + no bridge"]
    colors = [(119, 119, 119), (23, 63, 95), (0, 124, 131), (214, 107, 44), (163, 63, 123)]
    image = Image.new("RGB", (1200, 610), "white")
    draw = ImageDraw.Draw(image)
    draw.text((360, 18), "Exploratory simulation checks for the 40 °C digitized traces", fill="black")
    chart_top, chart_bottom = 90, 510
    for panel, concentration in enumerate(CONCENTRATIONS):
        left = 90 + panel * 570
        right = left + 470
        draw.text((left + 195, 58), f"{concentration} mg/mL", fill="black")
        for tick in (0, 25, 50, 75, 100):
            ytick = chart_bottom - int((chart_bottom - chart_top) * tick / 100)
            draw.line((left, ytick, right, ytick), fill=(220, 220, 220), width=1)
            draw.text((left - 35, ytick - 7), str(tick), fill="black")
        rows_c = [r for r in rows if int(r["concentration_mg_ml"]) == concentration]
        order = ["matched one-phase null", "matched two-phase alternative", "digitization perturbation"]
        if concentration == 180:
            order += ["180 offset + bridge sensitivity", "180 offset + bridge omitted"]
        values = [float(next(r for r in rows_c if r["case"] == c)["selected_k2_pct"]) for c in order]
        bar_w = 70
        gap = (right - left - len(values) * bar_w) / (len(values) + 1)
        for i, value in enumerate(values):
            x0 = int(left + gap + i * (bar_w + gap))
            x1 = x0 + bar_w
            y1 = chart_bottom - int((chart_bottom - chart_top) * value / 100)
            draw.rectangle((x0, y1, x1, chart_bottom), fill=colors[i], outline="black")
            draw.text((x0 + 10, y1 - 20), f"{value:.0f}%", fill="black")
            parts = labels[i].split()
            draw.text((x0 - 3, chart_bottom + 12), " ".join(parts[:2]), fill="black")
            if len(parts) > 2:
                draw.text((x0 - 3, chart_bottom + 27), " ".join(parts[2:]), fill="black")
    draw.text((20, 250), "K = 2 (%)", fill="black")
    image.save(OUT_PNG)

    print(f"Wrote {OUT_CSV}")
    print(f"Wrote {OUT_PNG}")
    for row in rows:
        print(row["concentration_mg_ml"], row["case"],
              f"K2={float(row['selected_k2_pct']):.1f}%",
              f"dAICc={float(row['median_delta_aicc']):.1f}")


if __name__ == "__main__":
    main()
```
