## Supplementary material for "Blind Recognition Reveals Early Multiphase IL-1ra Aggregation": Data/code package: run_simulation_validation.py

```
"""Operating-characteristic and perturbation checks for the blind hierarchy."""

from pathlib import Path
import csv
import sys

import numpy as np
from reportlab.lib.colors import Color, HexColor
from reportlab.pdfgen import canvas
import pypdfium2 as pdfium

HERE = Path(__file__).resolve().parent
ROOT = HERE.parents[1]
sys.path.insert(0, str(ROOT / "work" / "slide4_validation"))
from analyze_slide4 import fit_model, gompertz

OUT = ROOT / "outputs"
OUT_CSV = OUT / "simulation_validation_summary.csv"
OUT_PDF = OUT / "figs_v2" / "fig6_simulation_validation.pdf"
OUT_PNG = OUT / "figs_v2" / "fig6_simulation_validation.png"

T = np.arange(0.0, 6500.1, 50.0)
N_REP = 100
NOISE_SD = 0.008
PERTURB_SD = 0.010

# Parameters are baseline, then repeated (amplitude, inflection time, width),
# all in the same seconds/OD coordinates used by the manuscript fitter.
CASES = {
    "one phase": ([0.02, 1.9, 3000.0, 430.0], 1),
    "two phase, separated": ([0.02, 1.25, 2450.0, 260.0, 0.65, 4650.0, 520.0], 2),
    "two phase, overlapping": ([0.02, 1.35, 2850.0, 480.0, 0.65, 3700.0, 600.0], 2),
    "two phase, sub-threshold": ([0.02, 1.80, 2600.0, 360.0, 0.14, 4700.0, 460.0], 2),
}


def selection(fit1, fit2):
    p = fit2[2]
    a1, a2 = float(p[1]), float(p[4])
    fraction = min(a1, a2) / max(a1 + a2, 1e-12)
    gap = float(p[5] - p[2])
    width1 = float(p[3])
    delta = float(fit2[1] - fit1[1])
    accepted = delta < -10 and fraction >= 0.10 and gap > 2 * width1
    return accepted, delta, fraction, gap, width1


def fit_once(y):
    f1 = fit_model(T, y, 1)
    f2 = fit_model(T, y, 2)
    return selection(f1, f2)


def true_curve(params, k):
    return gompertz(T, params, k)


def load_digitized():
    data = {}
    with (OUT / "IL1ra_40C_digitized_traces.csv").open() as f:
        for row in csv.DictReader(f):
            c = int(row["concentration_mg_ml"])
            if c in (180, 200):
                data.setdefault(c, [[], []])
                data[c][0].append(float(row["time_s"]))
                data[c][1].append(float(row["od_digitized_corrected"]))
    return {c: (np.asarray(v[0]), np.asarray(v[1])) for c, v in data.items()}


def run():
    rng = np.random.default_rng(240731)
    rows = []
    examples = {}
    for name, (params, true_k) in CASES.items():
        y0 = true_curve(params, true_k)
        selected = []
        deltas = []
        fractions = []
        for rep in range(N_REP):
            y = y0 + rng.normal(0.0, NOISE_SD, size=T.size)
            result = fit_once(y)
            selected.append(2 if result[0] else 1)
            deltas.append(result[1]); fractions.append(result[2])
            if rep == 0:
                examples[name] = (y, result)
        rows.append({
            "dataset": name, "known_phase_count": true_k, "replicates": N_REP,
            "noise_sd_od": NOISE_SD,
            "selected_k1_fraction": selected.count(1) / N_REP,
            "selected_k2_fraction": selected.count(2) / N_REP,
            "median_delta_aicc_2_minus_1": float(np.median(deltas)),
            "median_candidate_smallest_amplitude_fraction": float(np.median(fractions)),
            "interpretation": "expected two-phase retention" if name == "two phase, separated" else
                              ("expected one-phase retention" if true_k == 1 else
                               "conservative rejection of collapsed/sub-threshold split"),
        })

    digitized = load_digitized()
    for c, (t, y) in sorted(digitized.items()):
        selected = []
        deltas = []
        for _ in range(N_REP):
            result = fit_model(t, y + rng.normal(0.0, PERTURB_SD, size=y.size), 2)
            one = fit_model(t, y + rng.normal(0.0, 0.0, size=y.size), 1)
            # Refit both models to the identical perturbed realization.
            perturbed = y + rng.normal(0.0, PERTURB_SD, size=y.size)
            one = fit_model(t, perturbed, 1)
            two = fit_model(t, perturbed, 2)
            accepted, delta, *_ = selection(one, two)
            selected.append(2 if accepted else 1); deltas.append(delta)
        rows.append({
            "dataset": f"digitized {c} mg/mL",
            "known_phase_count": "not known",
            "replicates": N_REP,
            "noise_sd_od": PERTURB_SD,
            "selected_k1_fraction": selected.count(1) / N_REP,
            "selected_k2_fraction": selected.count(2) / N_REP,
            "median_delta_aicc_2_minus_1": float(np.median(deltas)),
            "median_candidate_smallest_amplitude_fraction": "",
            "interpretation": "digitization-perturbation stability",
        })
    OUT_CSV.parent.mkdir(parents=True, exist_ok=True)
    with OUT_CSV.open("w", newline="") as f:
        writer = csv.DictWriter(f, fieldnames=list(rows[0]))
        writer.writeheader(); writer.writerows(rows)
    make_figure(examples)
    for row in rows:
        print(row["dataset"], row["selected_k2_fraction"], row["median_delta_aicc_2_minus_1"])


def panel(c, x, y, w, h, title):
    def xy(t, v): return x + t / 6500 * w, y + v / 2.25 * h
    for tx in (0, 2000, 4000, 6000):
        xx = xy(tx, 0)[0]; c.setStrokeColor(HexColor("#d9eafa")); c.setLineWidth(.4); c.line(xx, y, xx, y+h)
    for vy in (0, .5, 1, 1.5, 2):
        yy = xy(0, vy)[1]; c.setStrokeColor(HexColor("#d9eafa")); c.line(x, yy, x+w, yy)
    c.setStrokeColor(Color(.15,.15,.15)); c.setLineWidth(.7); c.line(x, y, x+w, y); c.line(x, y, x, y+h)
    c.setFillColor(Color(.1,.1,.1)); c.setFont("Helvetica-Bold", 7.5); c.drawString(x, y+h+6, title)
    return xy


def line(c, xy, ts, ys, color, width=1.0, dashed=False):
    c.setStrokeColor(HexColor(color)); c.setLineWidth(width); c.setDash(3,2) if dashed else c.setDash()
    p = c.beginPath()
    for i, (t, v) in enumerate(zip(ts, ys)):
        xx, yy = xy(t, v); p.moveTo(xx, yy) if i == 0 else p.lineTo(xx, yy)
    c.drawPath(p, stroke=1, fill=0); c.setDash()


def make_figure(examples):
    width, height = 540, 390
    c = canvas.Canvas(str(OUT_PDF), pagesize=(width, height))
    c.setFillColor(Color(.1,.1,.1)); c.setFont("Helvetica-Bold", 11)
    c.drawCentredString(width/2, height-18, "Simulation check of blind hierarchical phase selection")
    positions = [(54, 218), (285, 218), (54, 52), (285, 52)]
    dense = np.linspace(0, 6500, 500)
    for (title, (params, true_k)), (x, y) in zip(CASES.items(), positions):
        xy = panel(c, x, y, 190, 112, title)
        noisy, result = examples[title]
        fit1 = fit_model(T, noisy, 1); fit2 = fit_model(T, noisy, 2)
        line(c, xy, dense, gompertz(dense, fit1[2], 1), "#888888", .9, True)
        if result[0]:
            line(c, xy, dense, gompertz(dense, fit2[2], 2), "#173f5f", 1.4)
        else:
            line(c, xy, dense, gompertz(dense, fit1[2], 1), "#173f5f", 1.4)
        c.setFillColor(HexColor("#1f4e9a"))
        for t, v in zip(T, noisy):
            xx, yy = xy(t, v); c.circle(xx, yy, .65, stroke=0, fill=1)
        c.setFillColor(Color(.25,.25,.25)); c.setFont("Helvetica", 6.2)
        # Keep the diagnostic outside the data frame, on the panel title line.
        c.drawRightString(x+190, y+118, f"selected K={2 if result[0] else 1}; dAICc={result[1]:.1f}")
    c.setFillColor(Color(.2,.2,.2)); c.setFont("Helvetica", 7); c.drawCentredString(width/2, 17, "Time (s)")
    c.save()
    pdfium.PdfDocument(str(OUT_PDF))[0].render(scale=2.0).to_pil().save(OUT_PNG)


if __name__ == "__main__":
    run()
```
