## Supplementary material for "Blind Recognition Reveals Early Multiphase IL-1ra Aggregation": Data/code package: BDG_blind_early_multiphase.pdf

##### 2.3 Blind hierarchical phase recognition

The nested optical model is

$$g_K(t) = c + \sum_{j=1}^{K} a_j \exp\left[-\exp\left(-\frac{t-t_{0j}}{b_j}\right)\right], \qquad t_{01} < \cdots < t_{0K}, \tag{1}$$

where the peak optical rate of component $j$ is

$$R_{\mathrm{OD},j} = \frac{a_j}{eb_j}. \tag{2}$$

Model comparison used the Akaike Information Criterion corrected for small sample size (AICc), which scores a fitted model by combining its goodness of fit with a penalty for the number of free parameters [10]. Starting at $K = 1$, an added component was accepted only when $\Delta\mathrm{AICc} < -10$, every amplitude exceeded 10% of the total fitted amplitude, and every adjacent inflection-time gap exceeded two preceding-component time constants. In short, the recognition path is linear and stops at the first failure: $K = 1 \to$ fit $\to$ test $K+1$ against the three gates above $\to$ if all three pass, set $K = K+1$ and repeat; if any one fails, stop and retain the current $K \to$ continue until $K_{\max}$ is reached or a gate fails. The 53 °C and 40 °C series used $K_{\max} = 2$. The 50 °C series used $K_{\max} = 3$, ten independent starts, and at least 80% recurrence of the selected phase count. No observed rate, mass estimate, or response factor was supplied to this procedure.

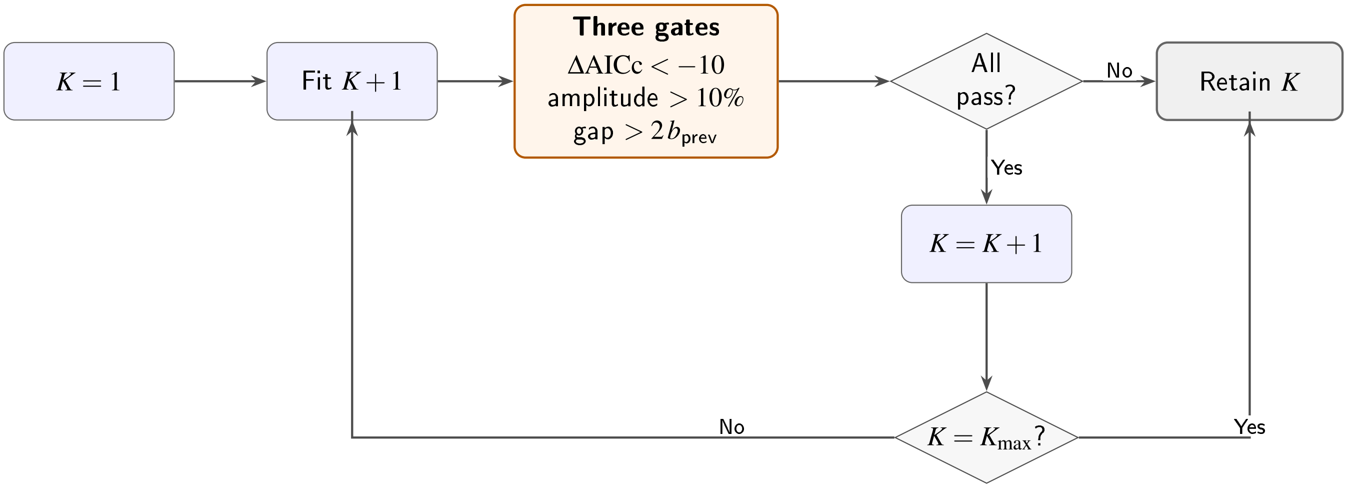


Figure 1: Blind hierarchical phase-recognition procedure. The loop starts by testing a two-phase fit against the one-phase baseline using the three gates. If any gate fails, one phase is retained and the procedure stops. If all three pass, two phases become the accepted result; the procedure then stops at two phases if $K_{\max}$ has been reached, or otherwise tests a three-phase fit against the two-phase baseline using the same gates, retaining three phases only if that fit also passes.

Assigning phase 1 to the high-response population and phase 2 to the low-response population gives $\alpha_1/\alpha_2 = 5.304$. For a two-phase trace, terminal mass-equivalent fractions are

$$F_1 = \frac{a_1/5.304}{a_1/5.304 + a_2}, \qquad F_2 = \frac{a_2}{a_1/5.304 + a_2}; \tag{3}$$

for a selected one-phase trace, $F_1 = 1$. With molecular weight 17.3 kDa, total protein concentration is $C_{tot} = 57.803p$ $\mu$M for $p$ in mg/mL. The conditional phase rate is

$$C_j = f_{agg} C_{tot} F_j, \qquad v_{j,\max} = \frac{C_j}{e b_j} \quad (\mu\text{M/min}). \tag{4}$$

Table 1: Synthetic and perturbation checks. Percentages are the fraction of 100 complete refits selecting the indicated phase count. Synthetic traces used 0.008 OD noise; digitized traces used 0.010 OD pointwise perturbations.

| Case | True $K$ | $N$ | Selected $K = 1$ % | Selected $K = 2$ % | Median ΔAICc (2–1) |
|---|---|---|---|---|---|
| One phase | 1 | 100 | 100 | 0 | 4.9 |
| Two phase, separated | 2 | 100 | 0 | 100 | -699.9 |
| Two phase, overlapping | 2 | 100 | 96 | 4 | -41.9 |
| Two phase, sub-threshold | 2 | 100 | 95 | 5 | -351.7 |
| Digitized 180 mg/mL | – | 100 | 0 | 100 | -221.1 |
| Digitized 200 mg/mL | – | 100 | 0 | 100 | -187.4 |

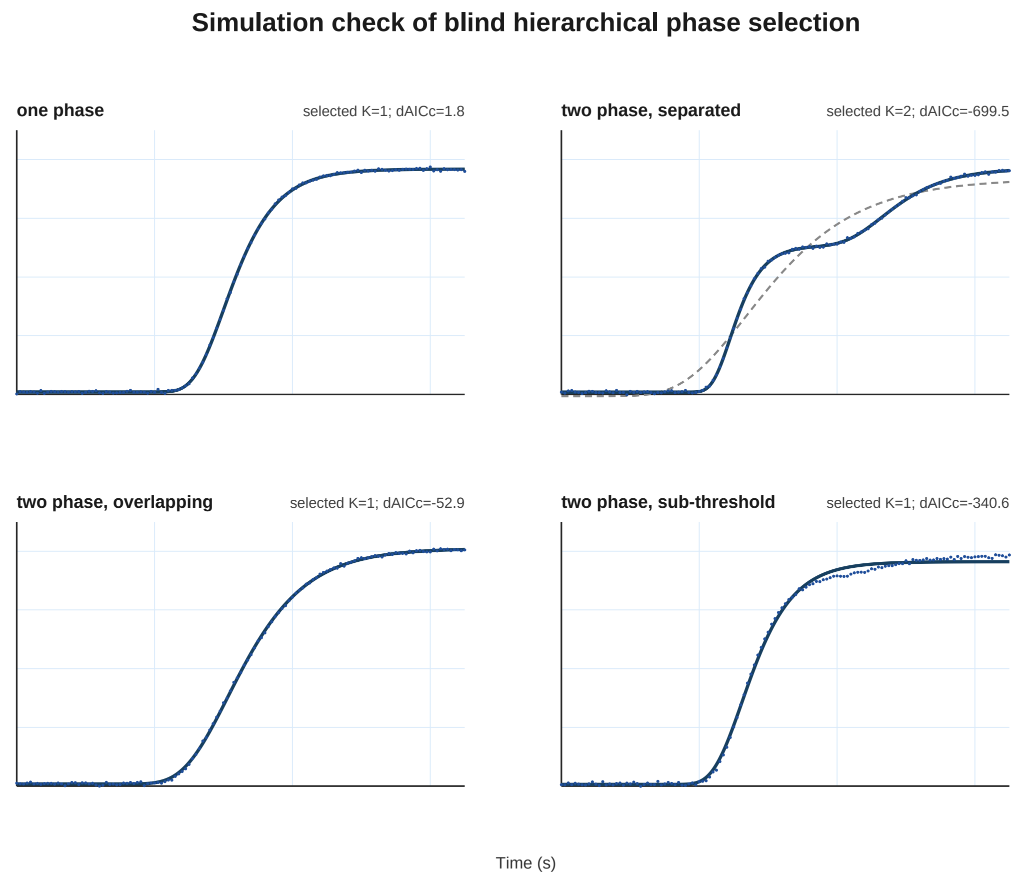


Figure 2: Simulation check of the blind hierarchy. Each panel shows one noisy realization with its one-phase fit (grey dashed) and the selected model (navy). The clearly separated two-phase trace is retained, whereas strongly overlapping and sub-threshold two-phase traces are conservatively retained as one phase. These simulations test operating characteristics and do not establish biological mechanism.

### 4 Results: 53 °C phosphate series

#### 4.1 Blind recognition resolves multiphase behavior at 5 mg/mL

One-phase fits appear visually strong at all four concentrations, but the blind hierarchy retained $K = 1$ at 1 and 2 mg/mL and selected $K = 2$ at 5 and 10 mg/mL (Fig. 3). At 5 and 10 mg/mL, the two-phase improvement was large (ΔAICc = -173.2 and -154.7, respectively). The candidate split at 1 mg/mL failed temporal separation, while that at 2 mg/mL lacked information support. The first resolved multiphase record is therefore 5 mg/mL, before the concentration range becomes high.

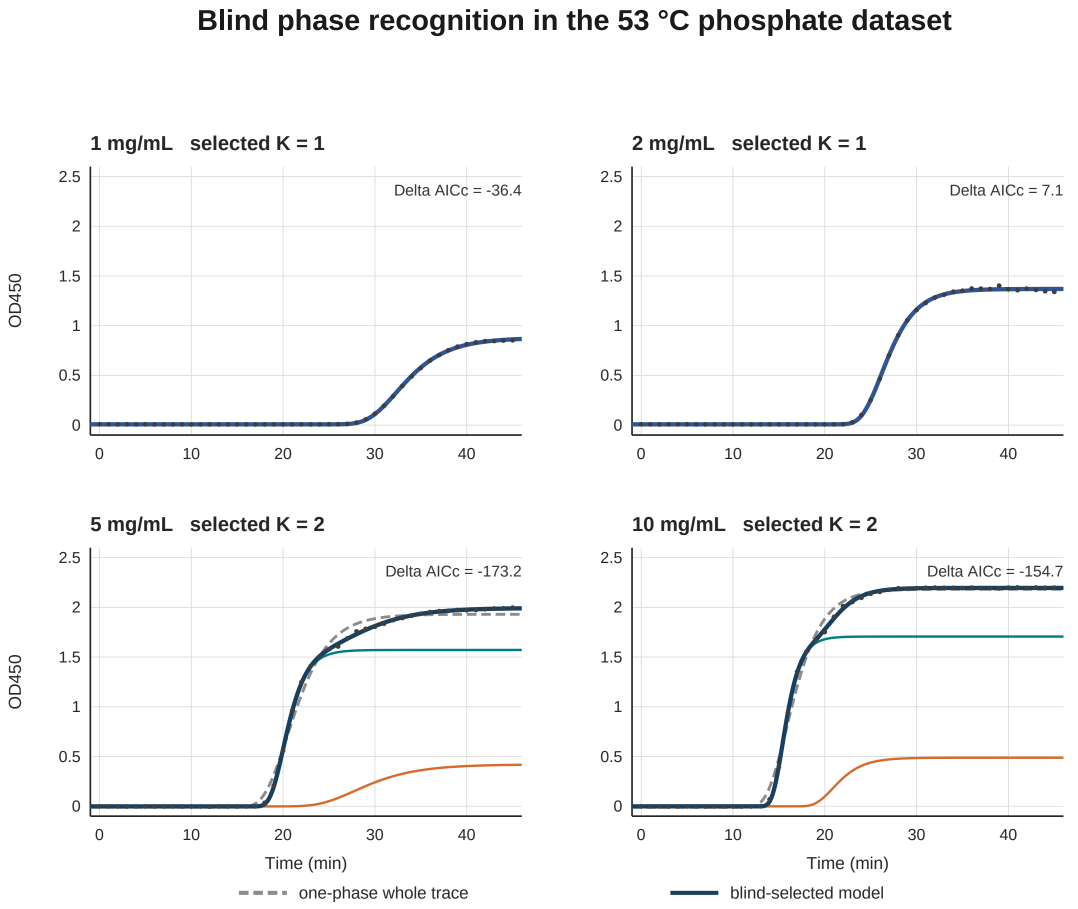


Figure 3: Blind phase recognition in the 53 °C phosphate series. Grey dashed curves are one-phase whole-trace fits; navy curves are blind-selected totals; teal and orange curves are the selected component contributions. The 1 and 2 mg/mL records remain one-phase; the 5 and 10 mg/mL records select two phases.

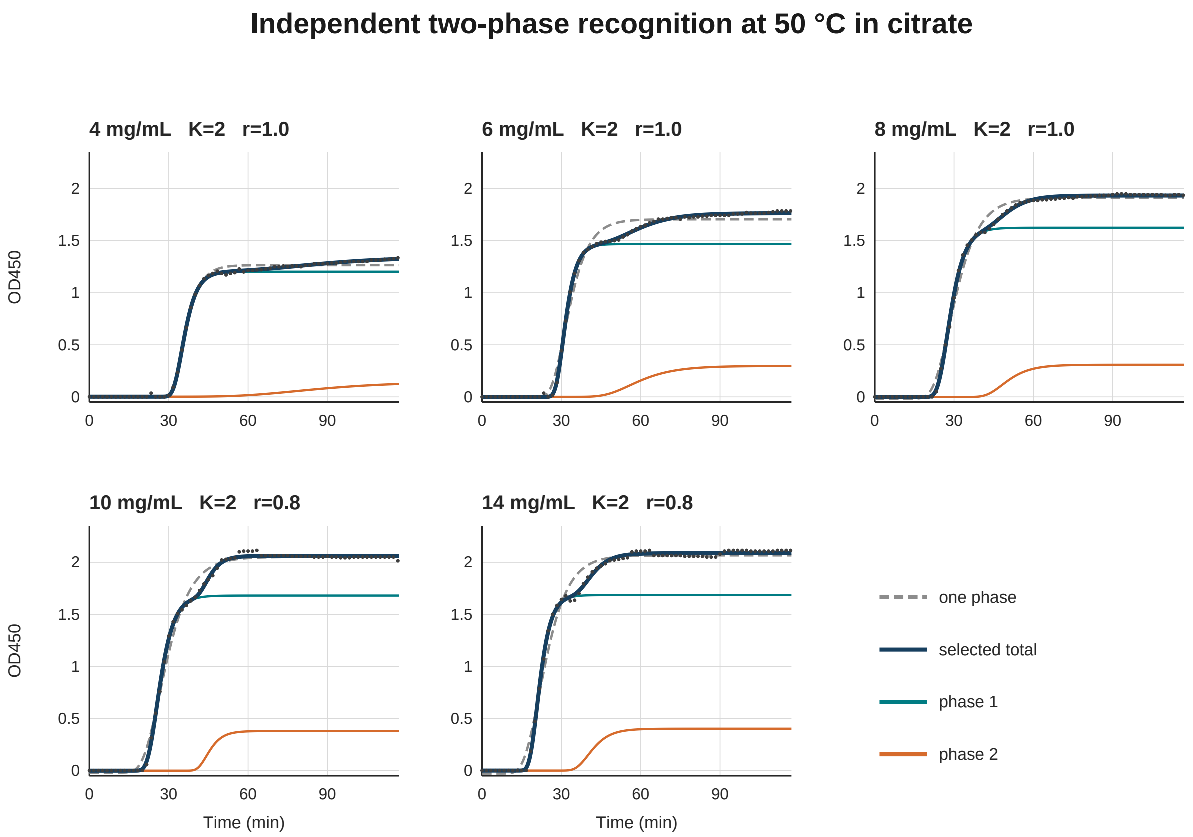


Figure 4: Compact redraw of the 50 °C citrate series. Points are digitized data; grey dashed curves are one-phase fits; navy curves are blind-selected totals; teal and orange curves are blind phase contributions. The 12 mg/mL record is excluded.

shoulder is not fitted perfectly by a single isolated component because both Gompertz terms contribute during the visible rise. Their overlap makes the individual onset parameters correlated, especially after digitization; the selected total, rather than either component alone, is the fitted observable. Thus the phase count is supported, whereas physical interpretation of a component's earliest tail should remain cautious.

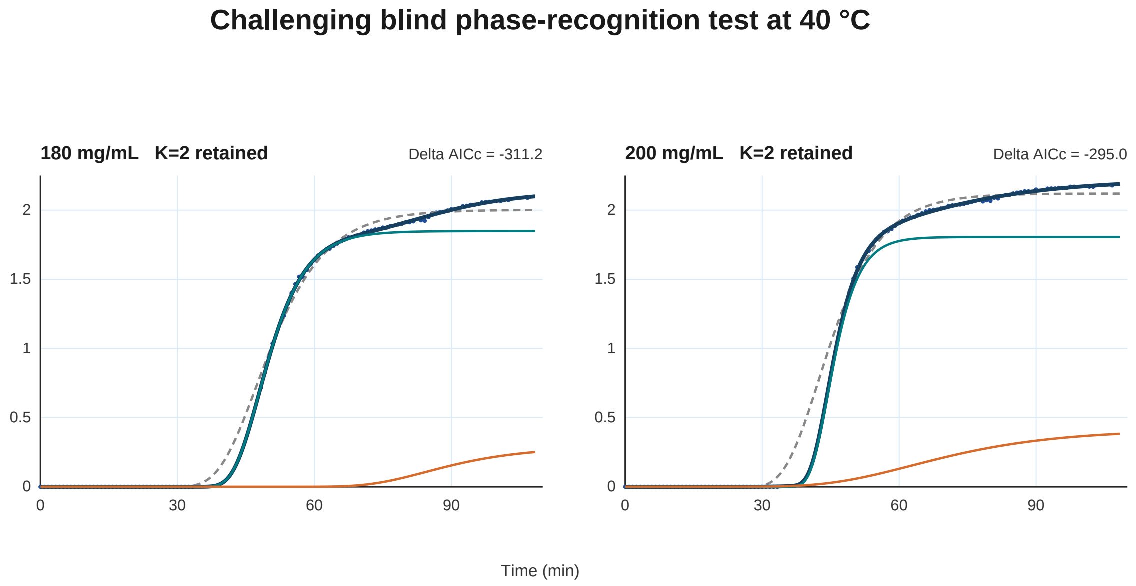


Figure 5: Challenging 40 °C high-concentration test. Points are reconstructed by digitizing the instrument-output plot at its displayed 50-s cadence. Grey dashed curves are one-phase fits; navy curves are blind-selected totals; teal and orange curves are the selected component contributions. The 180 mg/mL trace includes the documented correction of a visible instrument offset at approximately 5350 s. Both illustrative traces retain two phases. Component curves overlap during the leading rise, so the navy total is the relevant representation of fit quality.

`<username>/<repo>`. *[Placeholder: replace both URLs and the Zenodo DOI below once the repository and archive are created.]*
