## Supplementary material for "Blind Recognition Reveals Early Multiphase IL-1ra Aggregation": Data/code package: BDG_blind_early_multiphase_SI.tex

\documentclass[11pt]{article}
\usepackage[letterpaper,margin=0.76in]{geometry}
\usepackage[T1]{fontenc}
\usepackage{amsmath,amssymb}
\usepackage{mathptmx}
\usepackage{booktabs}
\usepackage{microtype}
\usepackage{hyperref}
\hypersetup{colorlinks=true,linkcolor=black,citecolor=black,urlcolor=black}
\setlength{\parskip}{0.40em}
\setlength{\parindent}{0pt}

\title{Supporting Information\\Blind Recognition Reveals Early Multiphase IL-1ra Aggregation}
\author{BDG Light --- Working Draft}
\date{31 July 2026}

\begin{document}
\maketitle

\section{Scope and data separation}

This supporting information reports the 53~$^\circ$C phosphate and 50~$^\circ$C citrate primary series separately, together with a separate 40~$^\circ$C high-concentration challenge set (180 and 200 mg/mL). Both primary series use pH 6.5, 140 mM NaCl, and 0.5 mM EDTA. The formulation-buffer component is 10 mM phosphate at 53~$^\circ$C and 10 mM citrate at 50~$^\circ$C; temperature and protein-concentration range also differ. The 40~$^\circ$C set is used for blind phase recognition only, not cross-series rate or mass comparison. No pooled fit, across-series score, or cross-series rate normalization is calculated. The 12 mg/mL citrate trace is omitted throughout.

\section{Blind selection rule}

For each trace, nested sums of ordered Gompertz components were fitted beginning at one component. A proposed additional component was retained only if: (i) \(\Delta\mathrm{AICc}<-10\); (ii) every component amplitude was greater than 10\% of total fitted amplitude; and (iii) every adjacent inflection-time gap exceeded two preceding-component time constants. The 53~$^\circ$C and 40~$^\circ$C series considered one or two components. The 50~$^\circ$C series considered up to three components using ten starts, with at least 80\% recurrence required for the selected count. Observed optical rates and the Type-I/Type-II conversion factor were excluded from this selection.

\section{Synthetic and sensitivity checks}

Four known-ground-truth traces were generated from the ordered Gompertz model: one phase, clearly separated two phases, strongly overlapping two phases, and a two-phase trace with a second amplitude below 10\%. Each trace contained 131 points at 50-s spacing. One hundred noisy realizations were fitted using 0.008 OD Gaussian noise and the same selection gates. For a 40~$^\circ$C-specific operating-characteristic check, 100 one-phase null traces were generated from each trace's fitted \(K=1\) curve and 100 two-phase alternatives from its fitted \(K=2\) curve, preserving the observed time points and adding 0.010 OD Gaussian noise. Separately, 0.010 OD Gaussian perturbations were added pointwise to each digitized 180 and 200 mg/mL trace for 100 complete refits. For 180 mg/mL, the post-offset correction was also drawn from \(0.0282\pm0.0050\) OD, with separate runs retaining or omitting the linear bridge point. These checks are sensitivity analyses within the fitted Gompertz model family, not estimates of instrument error or independent biological validation.

\begin{table}[htbp]
\centering
\small
\caption{Synthetic and digitization-perturbation checks.}
\begin{tabular}{lrrrr}
\toprule
Case & True $K$ & $K=1$ (\%) & $K=2$ (\%) & Median $\Delta$AICc\\
\midrule
One phase & 1 & 100 & 0 & 4.9\\
Two phase, separated & 2 & 0 & 100 & -699.9\\
Two phase, overlapping & 2 & 96 & 4 & -41.9\\
Two phase, sub-threshold & 2 & 95 & 5 & -351.7\\
Digitized 180 mg/mL & -- & 0 & 100 & -221.1\\
Digitized 200 mg/mL & -- & 0 & 100 & -187.4\\
\bottomrule
\end{tabular}
\end{table}

\newpage

\subsection{40~$^\circ$C-specific robustness check}

\begin{table}[htbp]
\centering
\small
\caption{40~$^\circ$C-specific matched simulation and correction-sensitivity checks. Each percentage is based on 100 complete refits. The matched null and alternative traces used the observed time points and 0.010 OD Gaussian noise. For 180 mg/mL, the offset-sensitivity runs varied the correction around 0.0282 OD and either retained or omitted the bridge point.}
\begin{tabular}{lrrrr}
\toprule
Case & 180 $K=2$ & 180 median & 200 $K=2$ & 200 median \\
 & (\%) & $\Delta$AICc & (\%) & $\Delta$AICc \\
\midrule
Matched one-phase null & 0 & 4.9 & 0 & 6.6 \\
Matched two-phase alternative & 100 & -255.5 & 99 & -217.4 \\
Digitization perturbation & 100 & -221.4 & 100 & -187.8 \\
180 offset + bridge sensitivity & 100 & -219.4 & -- & -- \\
180 offset + bridge omitted & 100 & -218.5 & -- & -- \\
\bottomrule
\end{tabular}
\end{table}

\begin{table}[htbp]
\centering
\small
\caption{Blind phase-selection summary for the 53~$^\circ$C phosphate series.}
\begin{tabular}{rrrrl}
\toprule
$p$ & selected $K$ & $\Delta$AICc for selected split & source marker & outcome\\
mg/mL & & & mOD/min & \\
\midrule
1  & 1 & split rejected & 104.6 & temporal separation failed\\
2  & 1 & split rejected & 237.4 & information support failed\\
5  & 2 & -173.2 & 408.0 & two phases retained\\
10 & 2 & -154.7 & 552.5 & two phases retained\\
\bottomrule
\end{tabular}
\end{table}

\section{40~$^\circ$C high-concentration challenge set}

The available 40~$^\circ$C record is a high-resolution SigmaPlot-style graphic derived from the instrument output, rather than a native numerical export. Marker coordinates for the retained 180 and 200 mg/mL traces were digitized at the displayed 50-s cadence. The 180 mg/mL trace contains a visible positive offset at approximately 5350 s; a 0.028 OD post-offset correction and a single linear bridge across the discontinuity were applied before fitting. The accompanying CSV identifies each baseline, digitized, corrected, and bridge value. This set is therefore a challenging test of phase recognition, not an independent raw-data validation. The individual empirical components overlap over the leading rise, so their earliest onset parameters are correlated; the selected total is the fitted observable. As a fit-quality diagnostic, the selected two-phase total reduced RMSE from 0.0368 to 0.0073 OD at 180 mg/mL and from 0.0293 to 0.0062 OD at 200 mg/mL relative to the one-phase fit. These residual reductions do not by themselves establish two biological mechanisms.

\begin{table}[htbp]
\centering
\small
\caption{Blind phase-selection summary for the 40~$^\circ$C high-concentration challenge.}
\begin{tabular}{rrrrrl}
\toprule
$p$ & selected $K$ & $\Delta$AICc & candidate P2 & P1--P2 gap & outcome\\
mg/mL & & (2--1) & amplitude fraction & s & \\
\midrule
180 & 2 & -311.2 & 0.135 & 2184 & two phases retained\\
200 & 2 & -295.0 & 0.187 & 1097 & two phases retained\\
\bottomrule
\end{tabular}
\end{table}

\begin{table}[htbp]
\centering
\small
\caption{Blind phase-selection summary for the 50~$^\circ$C citrate series. Printed Vmax values were not used in selection.}
\begin{tabular}{rrrrl}
\toprule
$p$ & selected $K$ & recurrence & printed Vmax & outcome\\
mg/mL & & starts/10 & mOD/min & \\
\midrule
4  & 2 & 10 & 108.34 & third phase unresolved\\
6  & 2 & 10 & 144.92 & third phase too small\\
8  & 2 & 10 & 160.97 & third phase too small\\
10 & 2 & 8  & 166.24 & third phase too small\\
14 & 2 & 8  & 175.19 & third phase too small\\
\bottomrule
\end{tabular}
\end{table}

\section{Observed-versus-blind rate comparisons}

The observed optical comparator is a withheld source marker in the 53~$^\circ$C series and a printed moving-window Vmax in the 50~$^\circ$C series. It is not a phase-specific observation and was not used by the blind procedure. Blind P1 and P2 are instantaneous Gompertz peak rates. The observed phase-1-equivalent \(\mu\mathrm{M}/\mathrm{min}\) value is obtained by multiplying the observed optical rate by the same-trace blind-P1 mass/optical ratio. It preserves the observed-versus-fit comparison after conversion but is not an independently measured mass rate.

\begin{table}[htbp]
\centering
\small
\caption{53~$^\circ$C phosphate series: observed and blind-fit rates before and after conversion. All mass-equivalent rates assume \(f_{\rm agg}=0.22\).}
\begin{tabular}{rrrrrrrr}
\toprule
$p$ & $K$ & observed & blind P1 & blind P2 & observed P1-equiv. & blind P1 & blind P2\\
mg/mL & & mOD/min & mOD/min & mOD/min & $\mu$M/min & $\mu$M/min & $\mu$M/min\\
\midrule
1  & 1 & 104.60 & 103.85 & --    & 1.531 & 1.520 & --\\
2  & 1 & 237.40 & 234.24 & --    & 4.434 & 4.375 & --\\
5  & 2 & 408.00 & 411.50 & 40.90 & 6.806 & 6.864 & 3.619\\
10 & 2 & 552.50 & 563.36 & 97.59 & 16.336 & 16.657 & 15.305\\
\bottomrule
\end{tabular}
\end{table}

\begin{table}[htbp]
\centering
\small
\caption{50~$^\circ$C citrate series: observed and blind-fit rates before and after conversion. All mass-equivalent rates assume \(f_{\rm agg}=0.22\).}
\begin{tabular}{rrrrrrrr}
\toprule
$p$ & $K$ & observed & blind P1 & blind P2 & observed P1-equiv. & blind P1 & blind P2\\
mg/mL & & mOD/min & mOD/min & mOD/min & $\mu$M/min & $\mu$M/min & $\mu$M/min\\
\midrule
4  & 2 & 108.34 & 138.23 & 2.46  & 2.808 & 3.583 & 0.338\\
6  & 2 & 144.92 & 192.34 & 11.67 & 3.637 & 4.827 & 1.553\\
8  & 2 & 160.97 & 163.26 & 19.22 & 5.014 & 5.085 & 3.176\\
10 & 2 & 166.24 & 172.58 & 42.11 & 5.710 & 5.927 & 7.671\\
14 & 2 & 175.19 & 214.27 & 32.31 & 8.166 & 9.988 & 7.988\\
\bottomrule
\end{tabular}
\end{table}

\section{Mass-equivalent calculation}

The Krishnan--Raibekas study \cite{KR} reports a 5.304-fold OD$_{450}$ response-per-mass ratio for Type I relative to Type II IL-1ra aggregate material. Blind phase 1 is assigned to the higher-response population and blind phase 2 to the lower-response population only for post-selection conversion. For a two-phase trace,
\begin{equation}
F_1=\frac{a_1/5.304}{a_1/5.304+a_2},\qquad
F_2=\frac{a_2}{a_1/5.304+a_2}.
\end{equation}
With molecular weight 17.3 kDa, \(C_{\rm tot}=57.803p\ \mu\mathrm{M}\) for \(p\) in mg/mL. The conditional peak mass-equivalent rate is
\begin{equation}
v_{j,\max}=\frac{0.22\,C_{\rm tot}F_j}{e b_j}\quad(\mu\mathrm{M}/\mathrm{min}).
\end{equation}
The sample volume is 0.2 mL, so the amount rate is \(0.2v_{j,\max}\) nmol/min. If the actual endpoint aggregate fraction is \(f\), all listed mass-equivalent rates scale by \(f/0.22\). The response factor and endpoint fraction are published external quantities, not measurements in either present kinetic series.

\begin{thebibliography}{99}
\bibitem{KR} S. Krishnan and A.A. Raibekas, ``Multistep Aggregation Pathway of Human Interleukin-1 Receptor Antagonist: Kinetic, Structural, and Morphological Characterization,'' \emph{Biophys. J.} 96, 199--208, 2009.
\end{thebibliography}

\end{document}
