## Supplementary material for "Blind Recognition Reveals Early Multiphase IL-1ra Aggregation": Data/code package: README.md

# Data and code package — Blind Recognition Reveals Early Multiphase IL-1ra Aggregation

Andrei A. Raibekas, Neurozon LLC. Companion package for the manuscript and its Supporting Information.

## Contents

- `manuscript/` — manuscript sources and compiled PDFs (main paper + Supporting Information).
- `figures/` — the four generated figures (fig1, fig3, fig5, fig6) as native vector PDFs, matching the versions embedded in the manuscript.
- `data/` — CSV tables underlying the paper's results:
  - `IL1ra_40C_digitized_traces.csv` — digitized 180/200 mg/mL, 40 °C instrument-output traces (baseline, digitized, offset-corrected, and bridge values), referenced in SI Section 4.
  - `observed_vs_blind_rate_comparison.csv` — observed vs. blind-fit phase rates for the 53 °C and 50 °C series, before and after mass-equivalent conversion (underlies SI Tables 6–7).
  - `phase_rates_both_temperature_sets.csv` — full per-trace fitted Gompertz parameters (amplitude, time constant, inflection time) and derived mass-equivalent quantities for both primary series.
  - `hierarchical_selection_summary_40C.csv` — AICc, amplitude fraction, and gap values behind the 40 °C two-phase selection (SI Table 4).
  - `simulation_validation_summary.csv` — operating-characteristic results for the four synthetic scenarios (SI Table 1 / Section 3).
  - `explore_40c_simulation_summary.csv` — matched-null and perturbation-sensitivity results specific to the 40 °C challenge (SI Section 3 / Table 2).
- `code/` — analysis scripts (Python) implementing the fitting and validation steps described in the Methods. See "Reproducibility notes" below before attempting to re-run them.

## Reproducibility notes — please read before re-running

This code was written for the author's own local project layout, not packaged as a standalone pipeline. Two limitations to know about:

1. **`analyze_slide4.py` and `digitize_40c_highres.py` require a raw source image that is not included in this package** (a screenshot/scan of the original instrument-output plot used for digitizing the 40 °C traces). Without that image, these two scripts cannot be re-run from scratch. The CSV they produce, `IL1ra_40C_digitized_traces.csv`, **is** included in `data/`, so every downstream step (fitting, validation) is reproducible starting from that file even without the raw image.
2. **`analyze_40c_challenge.py`, `explore_40c_simulation.py`, and `run_simulation_validation.py`** read the digitized CSV and reuse the `fit_model`/`gompertz` functions from `analyze_slide4.py`, but their file paths assume the author's original folder layout (e.g. `ROOT / "outputs" / ...`) rather than this package's `data/`/`code/` split. Expect to edit the `DATA`/`OUT` path constants near the top of each script to point at `../data/IL1ra_40C_digitized_traces.csv` (or copy the CSV alongside the script) before running.

None of this affects the manuscript's reported numbers — the CSVs in `data/` are the actual values used throughout the paper and SI tables, independent of whether the generating scripts are re-run.

## How to cite

Please cite the companion paper and, if using the tool specifically, the software record separately (see the manuscript's "Companion tool" section for the interactive tool's citation).

Raibekas, A.A. (2026). Blind Recognition Reveals Early Multiphase IL-1ra Aggregation. bioRxiv. [DOI pending]
