## Supplementary figures and images for "Blind Recognition Reveals Early Multiphase IL-1ra Aggregation"

### fig1_blind_phase_recognition_53C.pdf

#### Blind phase recognition in the 53 °C phosphate dataset

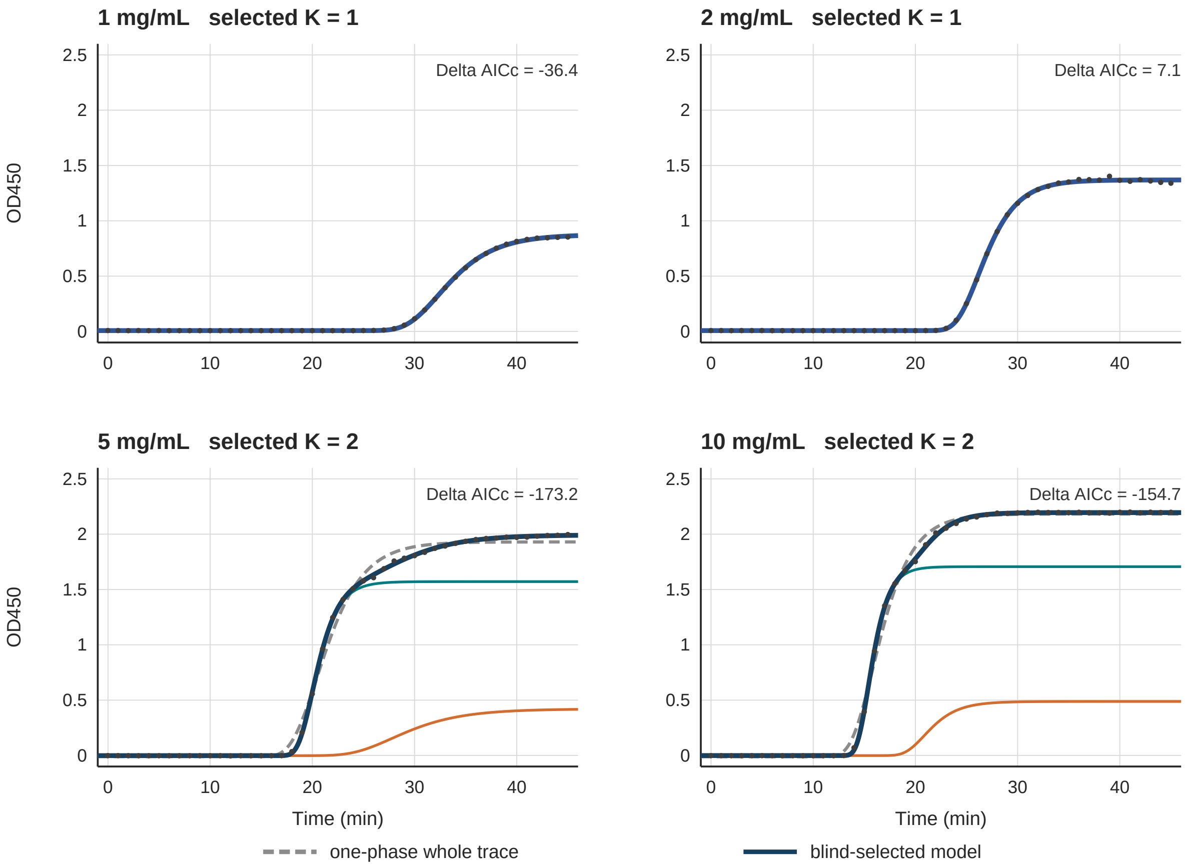

### fig3_compact_validation_50C.pdf

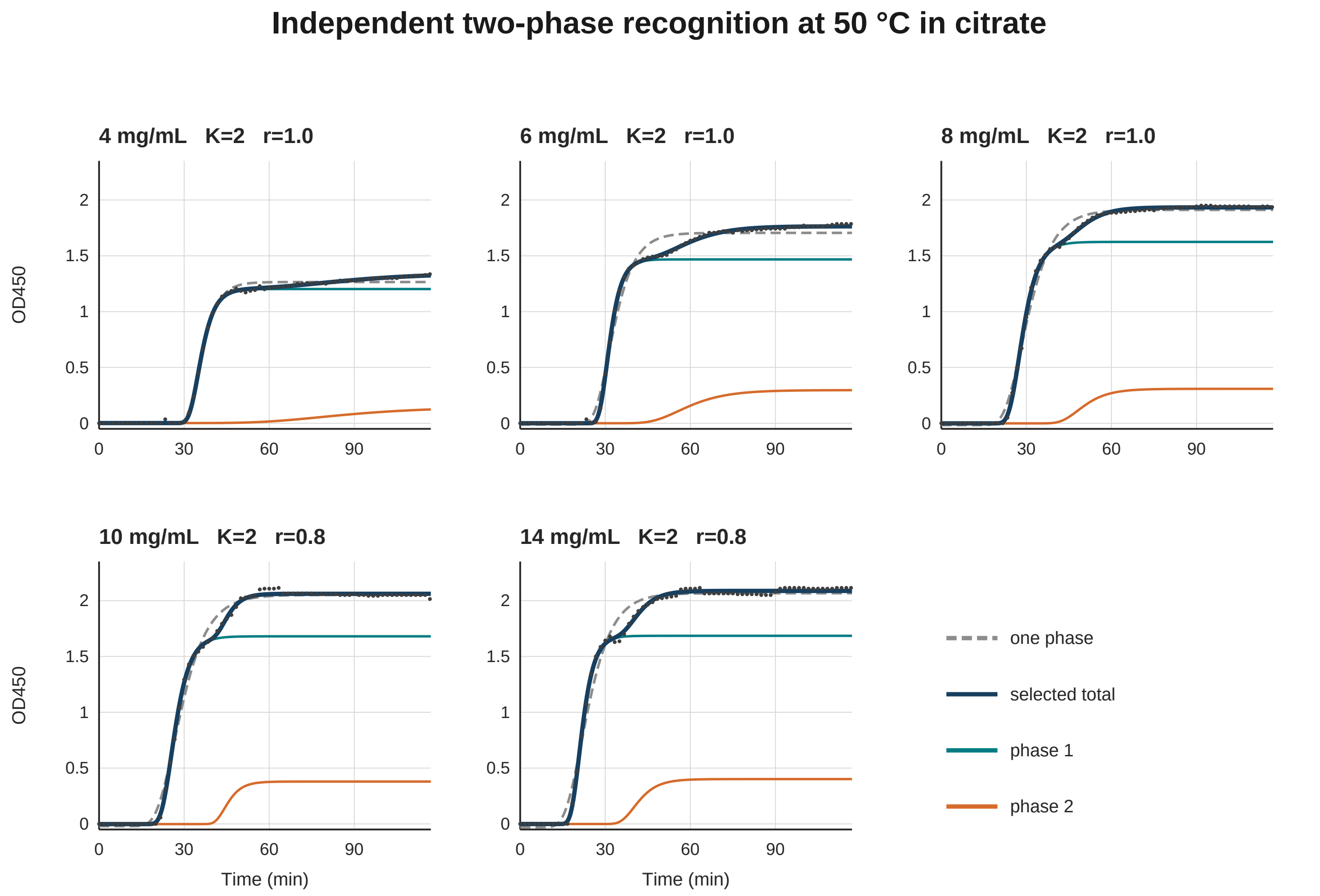

### fig5_40C_challenging_test.pdf

#### Challenging blind phase-recognition test at 40 °C

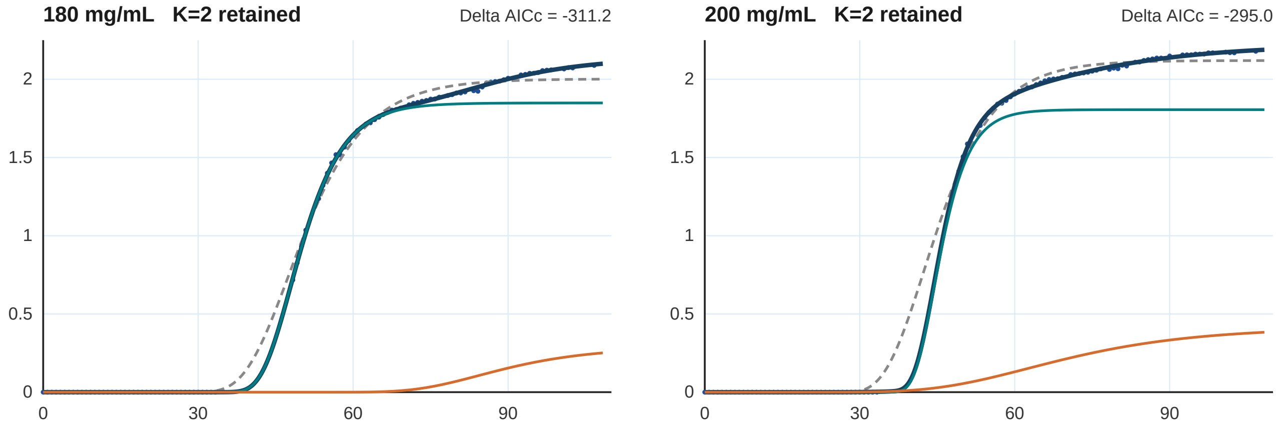


Time (min)

### fig6_simulation_validation.pdf

#### Simulation check of blind hierarchical phase selection

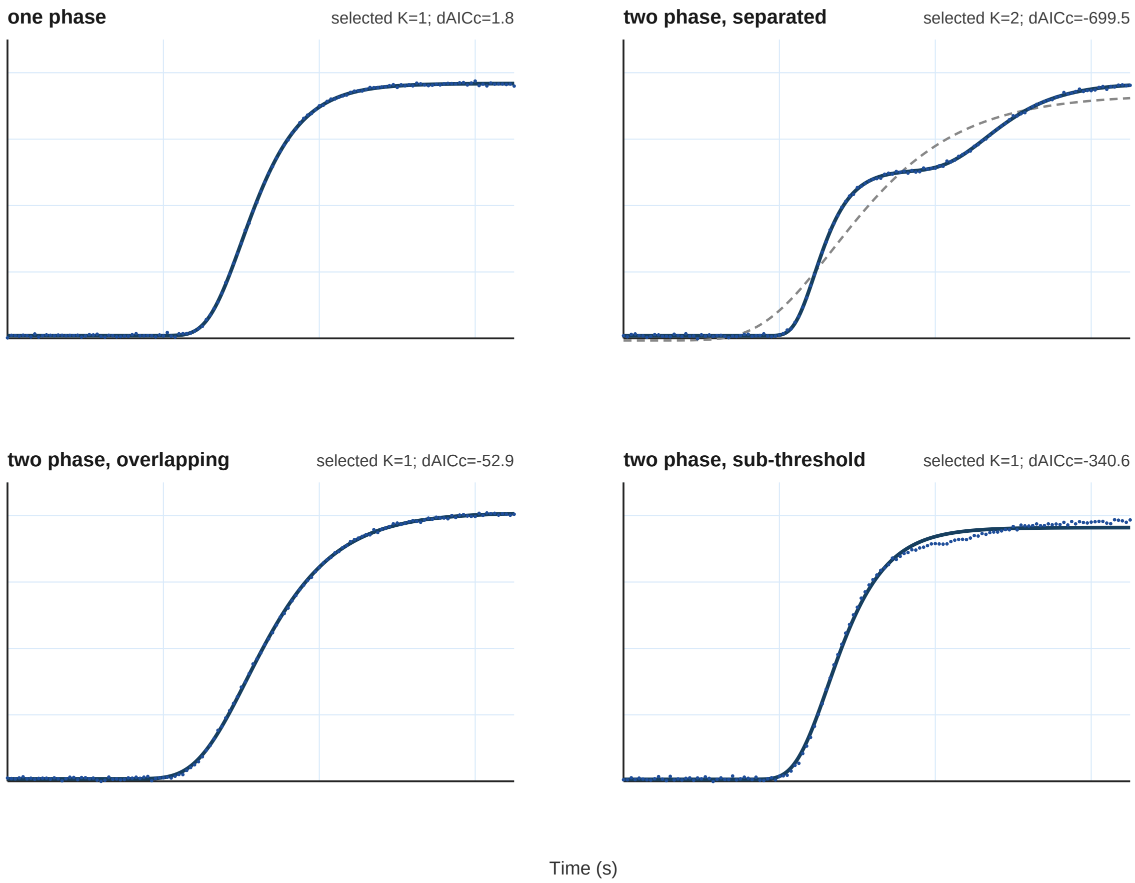
